# Conversion of an agonistic anti-TNFR2 biparatopic antibody into an antagonist by insertion of peptide linkers into the hinge region

**DOI:** 10.1101/2025.02.11.637430

**Authors:** Takuya Otsuki, Shigeyuki Matsumoto, Junso Fujita, Tomoko Miyata, Keiichi Namba, Ryo Kanada, Yasushi Okuno, Haruhiko Kamada, Hiroaki Ohno, Hiroki Akiba

**Author notes:** Corresponding authors, (H.A.), (H.O.).

## Abstract

Biparatopic antibodies (BpAbs) bind two different antigen epitopes to form characteristic immunocomplexes. Many BpAbs have been developed for enhanced cross-linking to induce signal transduction or cell internalization, whereas few were reported with smaller immunocomplexes to suppress unwanted signaling. Here, we developed a strategy to induce 1:1 immunocomplex formation to maximize antagonistic function. Various peptide linkers were introduced into the hinge regions of IgG-like agonist BpAbs against tumor necrosis factor receptor 2. Loss of crosslinking activity was observed for one BpAb, allowing the conversion of its function from an agonist to an antagonist. However, cross-linking activity was retained for another agonist BpAb, which binds to a different epitope pair. In a combined analysis of cryo-electron microscopy and coarse-grained molecular dynamics simulations, effect of epitope combination on the stability of 1:1 complexes was observed. These results lead to an understanding of the mechanism and design of BpAbs to adopt a 1:1-binding mode.

## Introduction

Antibodies efficiently bind to antigens based on their specificity, high affinity, and avidity (multivalency). Therapeutic antibodies utilize these characteristics. Conventional IgG antibodies are bivalent and contain two identical variable regions (Fvs) that recognize the same epitope on the surface of a specific antigen. This structural nature limits the immunocomplex to the antibody-to-antigen ratio of 1:2. However, the formation of a 1:2 complex is sometimes insufficient or may cause unwanted side effects, depending on the characteristics of the target molecule^1–3^. An attractive solution to this problem is the development of bispecific biparatopic antibodies (BpAbs)^4–5^. BpAbs are designed with two different Fvs binding to two different epitopes of the same antigen. This leads to the formation of diverse immunocomplexes other than the 1:2. BpAbs have been developed as IgG-like bivalent structure^6–10^ and tetravalent structures of IgG with additional antigen-binding domains^11–12^. Thus, an appropriate structure can be selected based on the required binding properties.

Tumor necrosis factor receptor superfamily (TNFRSF) proteins are potential targets for cancer immunotherapy, with advantages of using BpAbs. TNFRSF proteins are primarily involved in immunomodulation and apoptosis, and are activated by the formation of clusters on the cell membrane^13–15^. Antibodies that induce or suppress cluster formation can be used as agonists or antagonists, respectively. BpAbs are suitable for achieving these objectives by controlling the size of the immunocomplexes. Effective agonistic BpAbs against OX40 have been reported to induce cluster formation in cell membrane^16–17^. We also reported bivalent BpAbs targeting tumor necrosis factor receptor 2 (TNFR2) effectively producing both agonists and antagonists^18^. Furthermore, we recently reported successful design of an effective antagonist antibody against CD30 by combining epitopes based on the structural features of CD30^19^.

TNFR2 is primarily expressed on immune cells^20–23^. The extracellular region of TNFR2 comprises of four cysteine-rich domains (CRDs). By binding trimeric tumor necrosis factor α (TNFα) to CRD2 and CRD3, TNFR2 forms cluster on the cell membrane, and both classical and nonclassical NF-κB pathways are activated (**Figure 1a**) ^13, 24–27^. Since the expression of TNFR2 is associated with the proliferation of functional regulatory T cells, anti-TNFR2 antibodies can modulate immune functions (**Figure 1b**), and TNFR2 antagonists are expected to be used in cancer immunotherapy^28–31^. In our previous report^18^, bivalent BpAbs composed of five different Fvs binding different epitopes of TNFR2 were comprehensively produced. Effective agonists form clusters, whereas a potent TNFR2 antagonist exclusively forms a 1:1 complex. The relative position of the epitopes was considered the determinant of the activities of IgG-like BpAbs (**Figure 1c and 1d**)^18^. Since conventional IgG antibodies show moderate levels of agonistic activity owing to antibody:antigen = 1:2 complex formation, the bivalent BpAb format is crucial for producing a strong antagonist.

**Figure 1.**
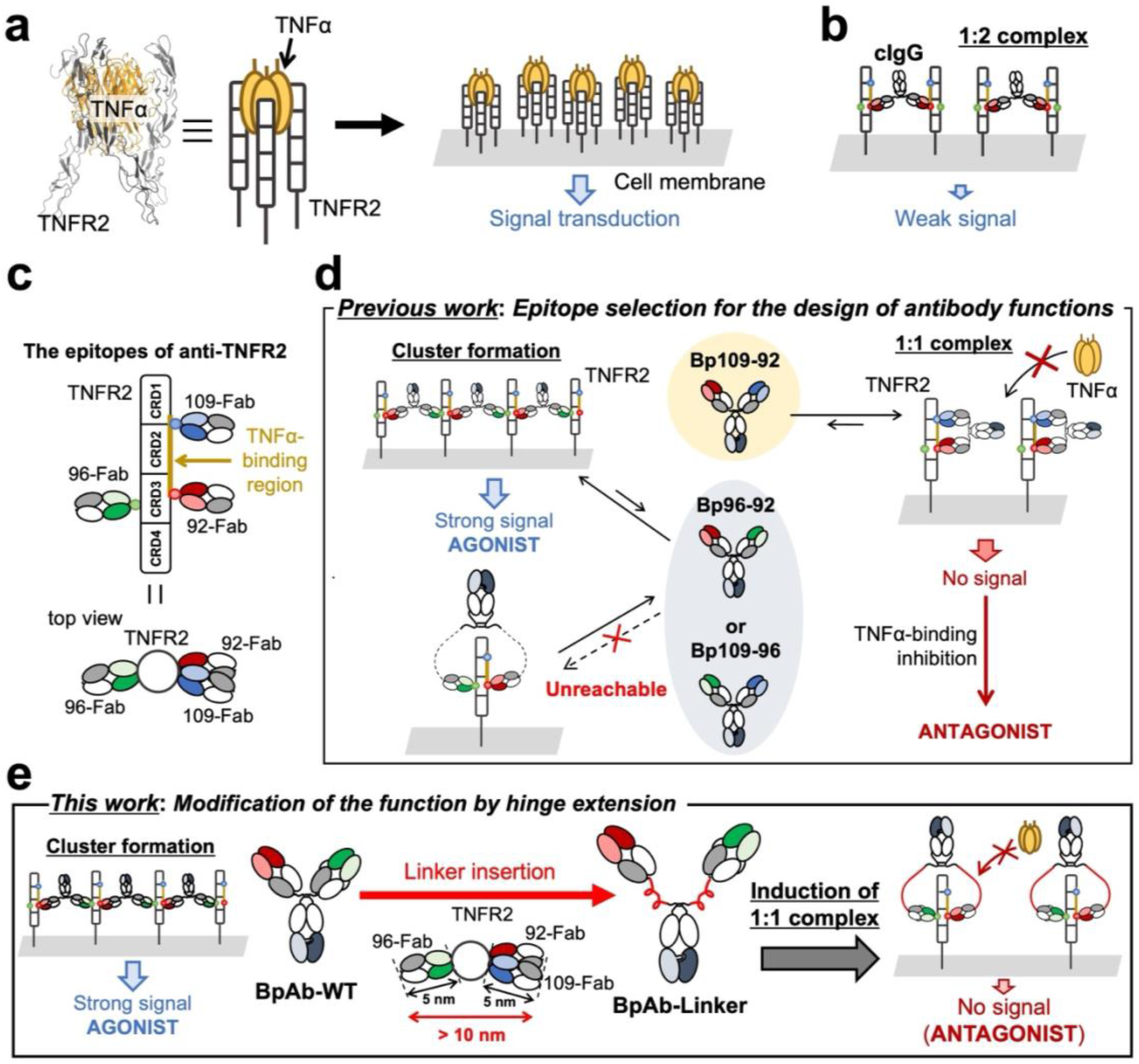
TNFR2 signaling and design of biparatopic antibodies (BpAbs). **a** Mechanisms of TNFR2 signaling. Trimer formation of TNFR2 on the cell membrane induced by the TNFα trimer triggers TNFR2 to form clusters and activate strong intracellular signaling. **b** Effect of conventional anti-TNFR2 antibody (cIgG) binding. CIgG crosslinks two TNFR2 molecules on the cell membrane to transduce a weak signal into the cell. **c** Epitope positions of the antibodies used for BpAb production in this study. 109-Fab and 92-Fab are competitive to TNFα. **d** Epitope selection for the design of IgG-like BpAbs reported previously^18^. Agonistic (Bp96-92 or Bp109-96) or antagonistic (Bp109-92) activities of BpAbs with conventional IgG structure is determined by the relative position of the pair of epitopes of the anti-TNFR2 monoclonal antibodies. **e** Design strategy of BpAb functions based on linker modification in this study. Linker insertion into the hinge region of an agonistic BpAb results in the induction of 1:1 complex formation to function as an antagonist.

In the previous study, agonistic BpAbs were produced from two Fvs binding distal epitopes in the ‘top view’ (**Figure 1c, bottom**). The distance between the C-termini of the two Fabs of IgG-like BpAbs in their bound state is the key to inducing cluster-forming abilities. The antigen-binding fragments (Fabs) of BpAbs are approximately 50 kDa, and the distance between the N– and C-termini is approximately 7 nm. For human IgG1, the two Fabs are linked by a short hinge region of approximately 2 nm. The hinge region is obviously too short to connect >10 nm distances between the C-terminal amino acids of two Fabs (92-Fab and 96-Fab or 109-Fab and 96-Fab) that bind distal epitopes on the surface of the same TNFR2 molecule. This hindered the formation of a 1:1 complex; alternatively, the formation of a large complex was favorable. Recently, IgG subclasses with different hinge structures have been reported to affect their biological activity based on their conformational degrees of freedom in immunocomplexes^32–35^. It has also been reported that the linker structure affects the activity of biparatopic DARPins^36–37^. Inspired by these observations, we hypothesized that the activity of anti-TNFR2 agonistic BpAbs is associated with the hinge structure and that they can be converted into non-agonists by inducing the formation of 1:1 complexes through the introduction of extended polypeptides (**Figure 1e**). In this study, we screened several peptides to extend the hinge regions of two agonistic BpAbs. We succeeded in suppressing the agonistic function of one BpAb and inducing an antagonistic function; thus, the same Fv pair could be used to switch between opposite functions depending on the linker structure. By analyzing the mechanisms of size-limited complex formation, we gained insights into the design of functional BpAbs.

## Results

### Biological activities of BpAbs inserted with various peptide linkers

We focused on three BpAbs designed by combining three Fvs (TR92, TR96, and TR109). TR92 and TR109 recognize the epitopes overlapping TNFα-binding region and they are competitive to TNFα. In a previous report, three BpAbs were developed in conventional IgG-like structures: two were strong agonists, whereas the other was a strong antagonist^18^. We first redesigned these anti-TNFR2 BpAbs into the CrossMab format^38^ because of difficulties in introducing linkers into published structures using intein-mediated protein trans-splicing (**Supplementary Figure 1**)^39^. BA1, BA2, and BA3 have the same Fvs as Bp96-92 (agonist), Bp109-96 (agonist), and Bp109-92 (antagonist), respectively, in the literature^18^. Two agonistic BpAbs (BA1-WT and BA2-WT) and one antagonist (BA3-WT) with the wild-type hinge sequence were obtained. Next, GP, G, P, AP10, EAAAK4, or Pro10 linkers were introduced into one of the heavy chains of the three BpAbs (**Table 1**, **Supplementary Tables 1-3** and **Supplementary Figures 2-4**)^40–44^. BA2-AP10 degraded too quickly to be evaluated (**Supplementary Figure 5a, lane 12**). Among the linkers used, GP, P, and Pro10 were relatively stable (**Supplementary Figure 6**). The signal-inducing activity of the BpAbs was evaluated using a reporter gene assay (**Supplementary Figure 7**). In the absence of the linker, the BpAbs showed activity similar to that of previously reported BpAbs^18^. The agonistic function of the linker-inserted BpAbs was not altered by any of the BpAb/peptide-linker combinations.

**Table 1.**
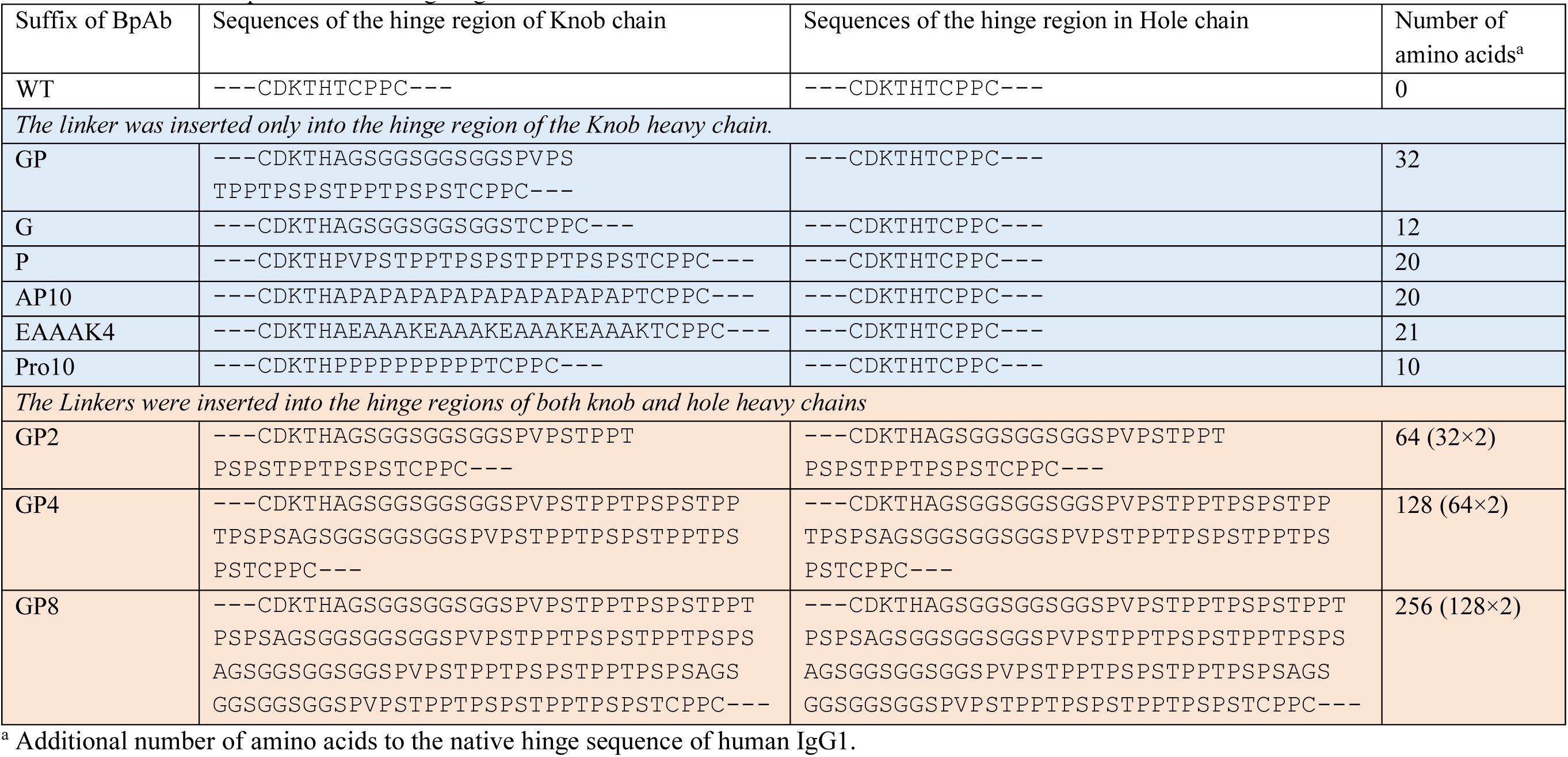
Amino acid sequences of the hinge region.

We assumed that the linkers were too short to observe any changes in activity. Therefore, further elongation of the linker was performed using the longest and most stable GP linker. One, two, or four units of the GP linker were inserted into both heavy chains of BA1-WT to produce BA1-GP2, BA1-GP4, or BA1-GP8 (**Figure 2a**, **Table 1**, **Supplementary Table 1** and **Supplementary Figures 5b**, **8**, **9**). BA2-WT was modified into BA2-GP2, BA2-GP4, and BA2-GP8 in a similar manner (**Supplementary Table 2** and **Supplementary Figures 5b, 10**, **11**). Signal-inducing and blocking activities were analyzed using reporter cells (**Figure 2b**). In the case of BA1, the maximum signal-inducing activity decreased with extending linker length, whereas the signal-blocking activity became more pronounced, especially for BA1-GP4 and BA1-GP8. A strong agonist, BA1-WT, without apparent signal-blocking activity, was converted into antagonists with minimal signal induction. The maximum signal-blocking activity was higher than those of the 92-Fab and 96-Fab mixtures (**Supplementary Figure 12**), which indicated the necessity of connecting two Fabs. In contrast, in the case of BA2, the maximum signal-inducing activities of BA2-GP2, BA2-GP4, and BA2-GP8 were reduced but only to half that of BA2-WT. The concentration dependency of the activity shifted towards higher concentrations upon linker insertion. The signal-blocking activity did not exceed that of the 109-Fab and 96-Fab mixtures (**Supplementary Figure 12**). In contrast to the agonists, the antagonist BA3-WT was not affected by the insertion of one or two units of the GP linker (**Supplementary Table 3** and **Supplementary Figure 5b, 13-15**).

**Figure 2.**
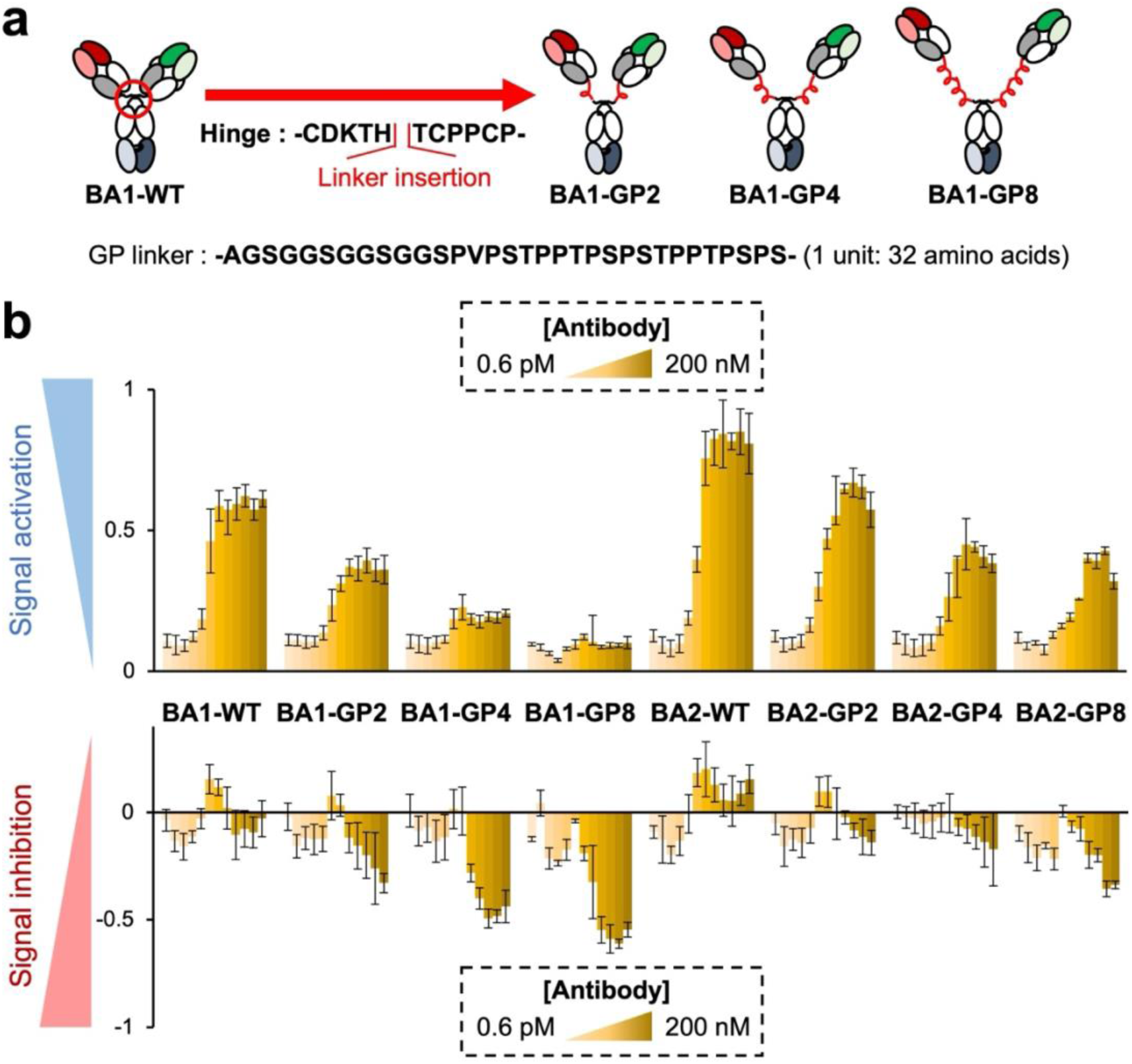
Effect of linker insertion in agonistic and antagonistic activities of the biparatopic antibodies (BpAbs). **a** Insertion of GP linkers into the hinge region in both of the two heavy chains of BA1-WT. BA2-WT was engineered in the same manner. **b** Agonistic (upper) and antagonistic (lower) activities of BpAbs after insertion of GP linkers in a reporter gene assay. Values are shown with the standard deviation of six independent experiments.

The GP linker used in this study was a peptide derived from the hinge of IgA^40^ and several Ser residues in this region were post-translationally modified with *O*-glycans^45^. The removal of *O*-glycans did not affect biological activities (**Supplementary Figure 16, 17**). Glycans were not important for conversion of agonistic BA1-WT into antagonistic BA1-GP4. Thus, linker length and epitope combination are considered primary factors for conversion.

### Apparent binding affinity of GP linker-inserted biparatopic antibodies

The apparent binding affinity of BpAbs to TNFR2-expressing reporter cells was investigated using flow cytometry (**Supplementary Figure 18**). The EC_50_ values of BA1-WT, BA1-GP2, and BA1-GP4 were 0.095, 0.097, and 0.107 nM, respectively. In the case of BA1, no change in the affinity was observed upon linker insertion. In contrast, the EC_50_ values for BA2-WT, BA2-GP2, and BA2-GP4 were 0.173, 0.219, and 0.475 nM, respectively, indicating that the apparent binding affinity decreased depending on linker length. This was consistent with the concentration dependency of BA2 in the reporter gene assay; by the insertion of a longer linker, higher concentrations were required to achieve maximum activity. However, the differences in maximal agonistic activity that are dependent on epitope selection are not understood.

### Analysis of the size of complexes between anti-TNFR2 biparatopic antibody and TNFR2

To evaluate the crosslinking binding pattern associated with signal transduction, the sizes of the immunocomplexes were analyzed. First, the mixture of BpAb and TNFR2 C-terminally fused with maltose binding protein (TNFR2-MBP) was mixed at 1 μM each in PBS and analyzed in size-exclusion chromatography (SEC) (**Figure 3a**). While elution volume of BA1-WT and TNFR2-MBP was 16.2 and 16.7 mL, and the elution volume ranged from 11 to 15 mL, with a maximum elution volume of 13.6 mL. A smaller elution volume indicates the formation of large immunocomplexes. The elution volume of BA1-GP2 was smaller than that of BA1-WT, reflecting a larger hydrodynamic radius^40^. However, the elution volume of the BA1-GP2–TNFR2-MBP mixture shifted to a larger volume than that of BA1-WT–TNFR2-MBP mixture, with a maximum of 14.6 mL. This indicated a tendency to favor smaller immunocomplexes in the case of BA1-GP2 than in BA1-WT. For BA1-GP4, elution of the mixture showed a dominant peak at 14.6 mL, with only 0.4 mL shift from BpAb alone (15.0 mL). Formation of a small, specific immunocomplex is indicated. The complex formed between BA1 and TNFR2 shrank in a linker length-dependent manner. These results are consistent with the reduction in signal-inducing activity by linker insertion in a reporter gene assay. For BA2-GP2 and BA2-GP4, larger BpAbs and smaller immunocomplexes were observed by linker insertion into BA2-WT. However, in the case of BA2-GP4, the dominant peak was not clear, and immunocomplexes of diverse sizes were observed for BA1-GP2 and BA2-GP2.

**Figure 3.**
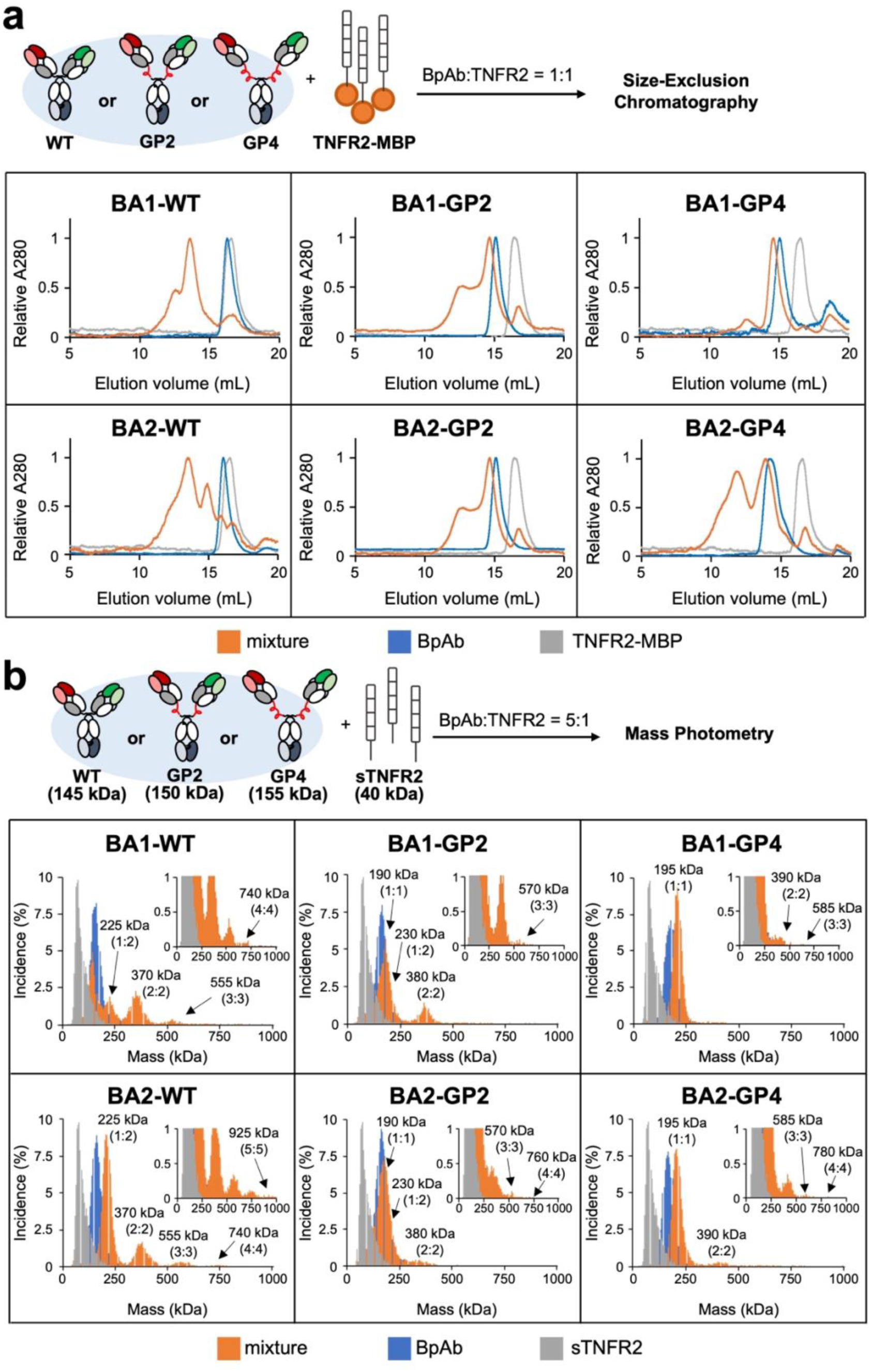
Immunocomplex formed between the biparatopic antibodies (BpAbs) and TNFR2. **a** Size-exclusion chromatography for the analysis of size distribution. **b** Mass photometry for the analysis of relative frequency of the observed immunocomplexes with their mass.

The size of the immunocomplexes was analyzed at a single-particle level for mixtures of BpAb (27.5 nM) and TNFR2 digested from etanercept (sTNFR2, 5.5 nM) using mass photometry (**Figure 3b**). First, the molecular weights of the individual proteins were measured and were found to be 145 kDa (BA1-WT and BA2-WT), 150 kDa (BA1-GP2 and BA2-GP2), 155 kDa (BA1-GP4 and BA2-GP4), and 40 kDa (sTNFR2), respectively. Next, the molecular weights of the complexes present in mixtures of BpAbs and sTNFR2 were measured. For BA1-WT and BA2-WT, peaks with molecular weights corresponding to antibody:antigen = 1:2 (225 kDa) and larger equimolar complexes (2:2 [370 kDa], 3:3 [555 kDa] and other larger particles) were observed. The large immunocomplexes observed in SEC were derived from these crosslinking complexes. In contrast, the 1:1 complex (190 or 195 kDa) was the major immunocomplex observed in the linker-inserted BpAbs. For BA1-GP2, the 2:2 complex (380 kDa) was observed at a slightly lower level than BA1-WT. A small amount of 3:3 complex (570 kDa) was also observed. Furthermore, only a small amount of the 2:2 complex (390 kDa) was observed in BA1-GP4, and the 1:1 complex was nearly dominant. In the cases of BA2-GP2 and BA2-GP4, only a small amount of the 2:2 complex was maintained. These observations are consistent with those of SEC analysis, although lower concentrations may have shifted the equilibrium to smaller immunocomplexes. The results of the size analysis of immunocomplexes clearly showed a correlation with the function; agonistic function was observed for BpAbs that showed the formation of 2:2 or larger immunocomplexes in solution. This observation is consistent with a previous study on epitope variations^18^. BA1-GP4 lost its agonistic activity upon linker insertion because of the favorable formation of a 1:1 complex, as expected.

BA3-WT, BA3-GP2, and BA3-GP4 complexed with sTNFR2 were analyzed using mass photometry. Exclusive formation of the 1:1 complex was maintained by linker insertion (**Supplementary Figure 19**), which was consistent with the reporter gene assay (**Supplementary Figure 15**).

### Structural analysis of 1:1 complex between BA1-GP4 and TNFR2 via cryo-electron microscopy

To further understand the mechanism of 1:1 complex formation, cryo-electron microscopy (EM) single-particle analysis was performed to determine the tertiary structure of the complex. The complex structure of BA1-GP4–TNFR2-MBP was determined at a resolution of 3.73 Å (**Figure 4a**, **Supplementary Table 4** and **Supplementary Figure 20**-21). Two Fabs (92-Fab and 96-Fab) of BA1-GP4 bound simultaneously to two epitopes of TNFR2. No densities corresponding to the linker, Fc, or MBP were observed. Most of the CRD4 in TNFR2 was not clearly observed. The absence of these regions, indicating flexibility, was similar to that previously observed for Bp109-92 (original to BA3-WT)^18^. The density corresponding to CRD1 of TNFR2 and the constant region of the two Fabs was partially low, but this did not affect the subsequent steps of epitope determination with the orientation of Fab bound to TNFR2.

**Figure 4.**
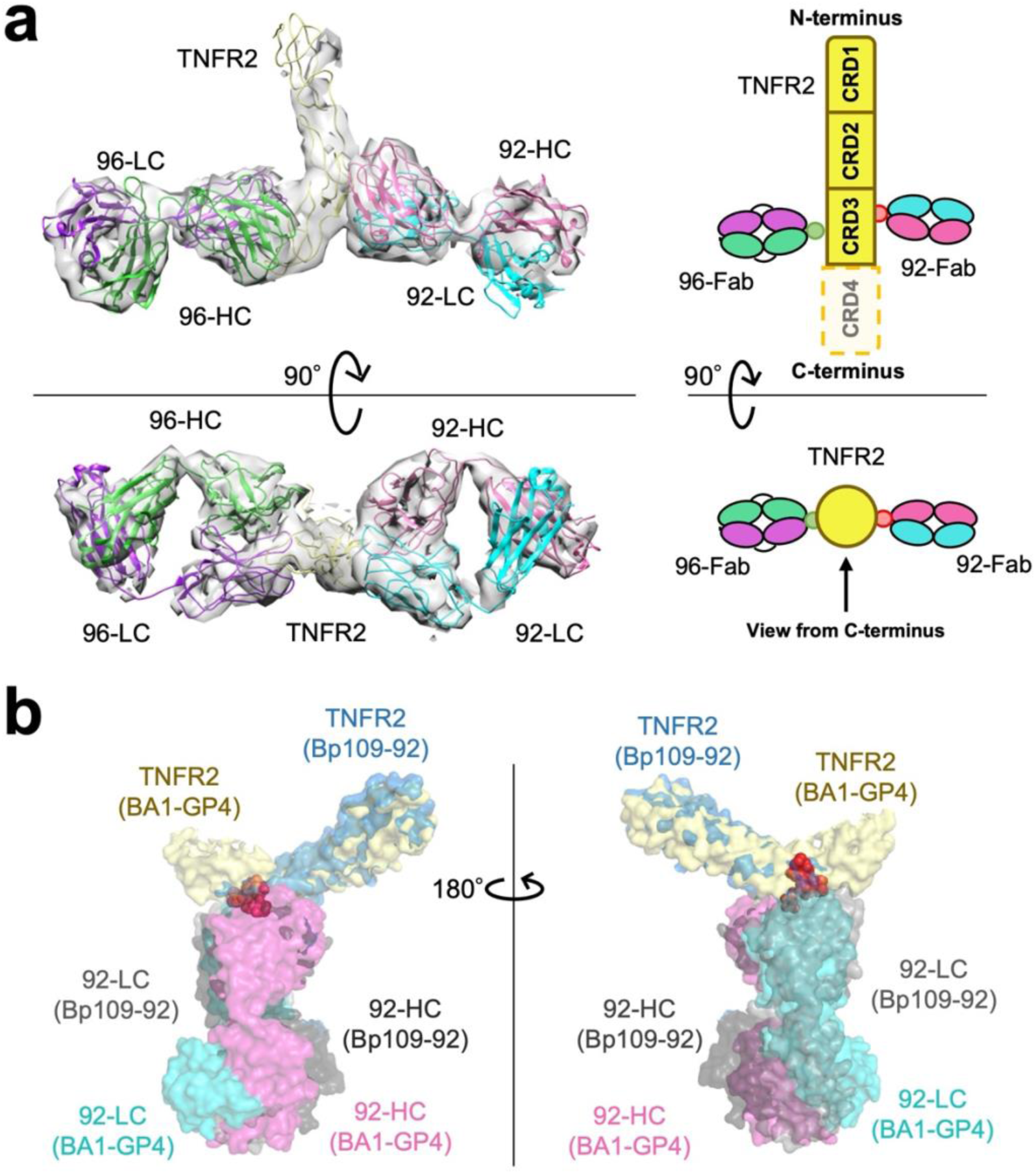
Structure of the BA1-GP4–TNFR2 complex. **a** Overall view of the cryo-EM structure of BA1-GP4–TNFR2 complex with a schematic diagram showing cysteine-rich domains (CRDs). Heavy and light chains from TR92 and TR96 are described as 92-HC (magenta), 92-LC (cyan), 96-HC (green) and 96-LC (purple), respectively. For TNFR2 (yellow), CRD4 was only partially observed (split lines). No density corresponding to the linker, Fc, and MBP was observed. The absence of these regions indicated the flexibility. **b** Binding interface of TR92-Fab–TNFR2 in two different biparatopic antibodies. BA1-GP4–TNFR2 (PDB ID: 9LFL) and Bp109-92–TNFR2 (PDB ID: 8HLB) complexes are aligned with the mutant-defined epitope residues of TR92 (red spheres; see **Supplementary Figure 22**).

The TNFR2-bound structure of 92-Fab has already been solved as part of the Bp109-92–TNFR2 complex^18^. The two TNFR2–92-Fab interfaces were analyzed using the PISA server and the epitope residues were almost unaffected (**Supplementary Figure 22**)^46^. When the coordinates were aligned using epitope residues on the TNFR2 side, the orientation of Fab to TNFR2 was unaffected (**Figure 4b**). Thus, the two structures using different BpAbs retained the molecular recognition of 92-Fab as their component. TNFR2-bound structure of 96-Fab was newly determined in this study. The epitope of this antibody was previously determined by mutagenesis^18^. This region was consistently identified as the epitope of the complex structure (**Supplementary Figure 23**).

Notably, 92-Fab and 96-Fab bound to TNFR2 at approximately the same position along the long axis of TNFR2, and their paratopes faced each other in a 180° orientation. Given this structural feature, we hypothesized that the BA1-GP4–TNFR2 1:1 complex may be formed without distorting the flexible conformation of TNFR2, whereas for BA2-GP4 possessing 96-Fab and 109-Fab, 1:1 complex formation could induce distortion of TNFR2, resulting in different behaviors in immunocomplex formation.

### Structural motions of TNFR2 in the 1:1 complexes with anti-TNFR2 BpAbs analyzed by coarse-grained molecular dynamics (CGMD) simulation

To understand the mechanism of epitope-dependent differences in the immunocomplex formation of BpAbs with TNFR2, structural stability of TNFR2 in the 1:1 complex was analyzed using coarse-grained molecular dynamics (CGMD) simulations with the program CafeMol^47^. The initial structures of the 1:1 complex of four BpAbs with extended linkers (BA1-GP2, BA1-GP4, BA2-GP2, or BA2-GP4) and TNFR2 were modeled using two PDB files (PDB ID: 8HLB, 9LFL; **Figure 5a** and **Supplementary Figure 24**-28).

**Figure 5.**
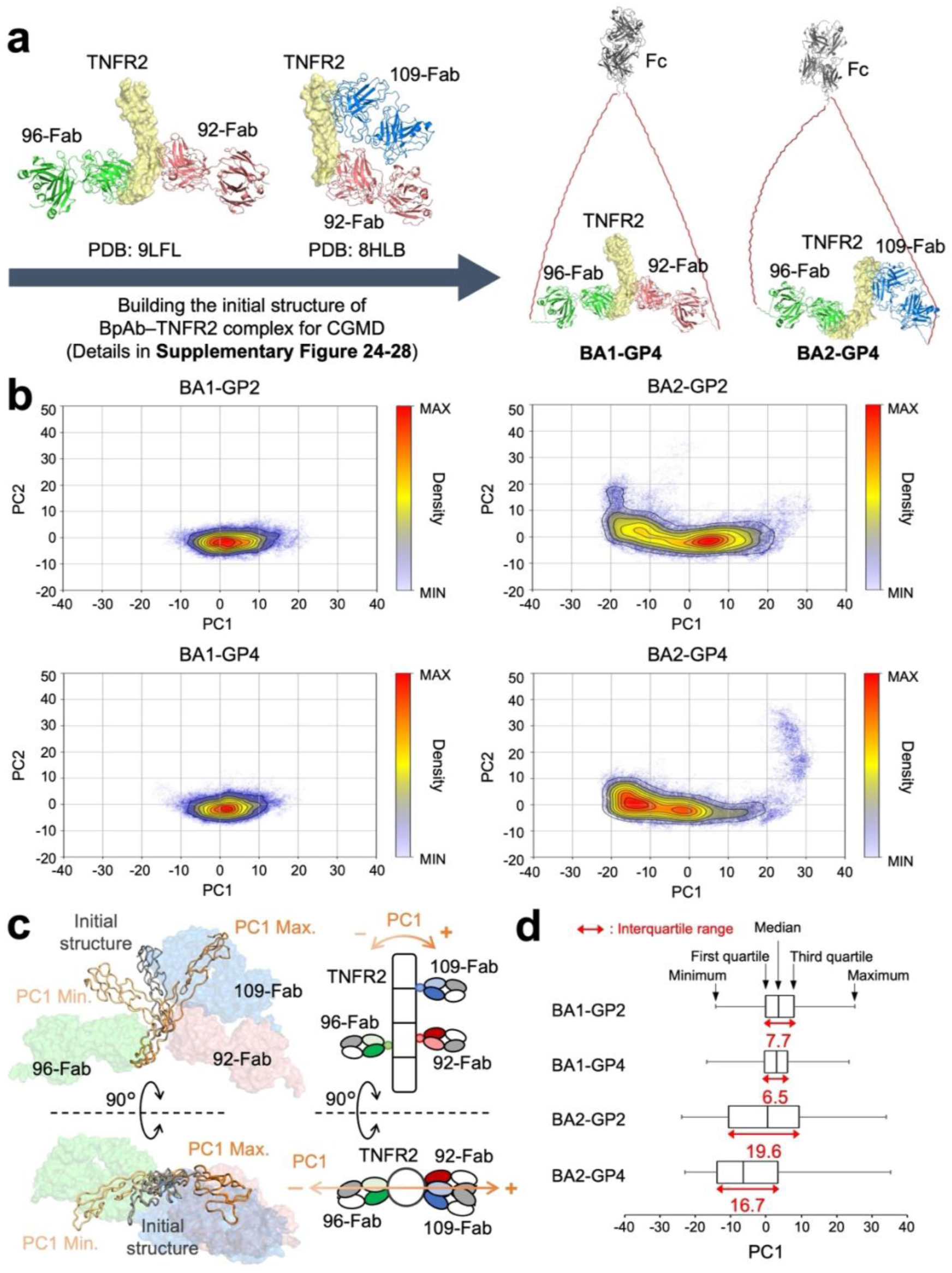
Coarse-grained molecular dynamics (CGMD) simulation for the analysis of structural features of TNFR2 in 1:1 immunocomplexes with BpAbs. **a** Initial structure of the complex between TNFR2 and BpAbs (BA1-GP2, BA1-GP4, BA2-GP2 and BA2-GP4) for CGMD (see also **Supplementary Figure 24**-28). **b** Principal component analysis (PCA) of the structures of TNFR2 in 10 runs of 10^6^ steps of the CGMD simulation shown with density-based clustering for each sample by the two main components. **c** Structural changes of TNFR2 along PC1-axis. **d** Distribution of the PC1 values.

Through the simulation runs, binding interfaces of the Fabs were all maintained for four BpAb– TNFR2 complexes. Differences were observed in the structures of TNFR2. Structural motions of TNFR2 in complex with BpAb was analyzed through principal component analysis using the coordinates of all Cα atoms of TNFR2 (derived from ten independent trajectories of 10^6^ steps in CGMD simulations). (**Figure 5b**). The contribution of the first principal component (PC1) was 65.7% (**Supplementary Table 5**), indicating that the dominant differences among runs with different BpAbs could be explained by this component. The movement along the PC1 axis coincided with the movement of the N-terminus of TNFR2 in the direction of Fab binding (**Figure 5c**). Compared with BA1-GP2, BA2-GP2 exhibited a greater structural change along the PC1 axis (**Figure 5d**). Similarly, BA2-GP4 induced a larger structural change along the PC1 axis compared to BA1-GP4 (**Figure 5d**). Independent of the linker length, larger structural motions of TNFR2 in complex with BA2 were observed compared to those in complex with BA1.

## Discussion

In this study, we investigated changes in the function of BpAbs by inserting various peptide linkers into the hinge region. The most important finding was the conversion of the agonist, BA1-WT into an antagonist by insertion of the long linkers of BA1-GP4 and BA1-GP8 (**Figure 2**). The original agonistic BpAb formed cross-linked complexes with TNFR2, but the insertion of four GP linkers resulted in the formation of a stable 1:1 complex with antagonistic function (**Figure 6a**). Notably, expanding the size of BpAbs led to shrinkage of the immunocomplex.

**Figure 6.**
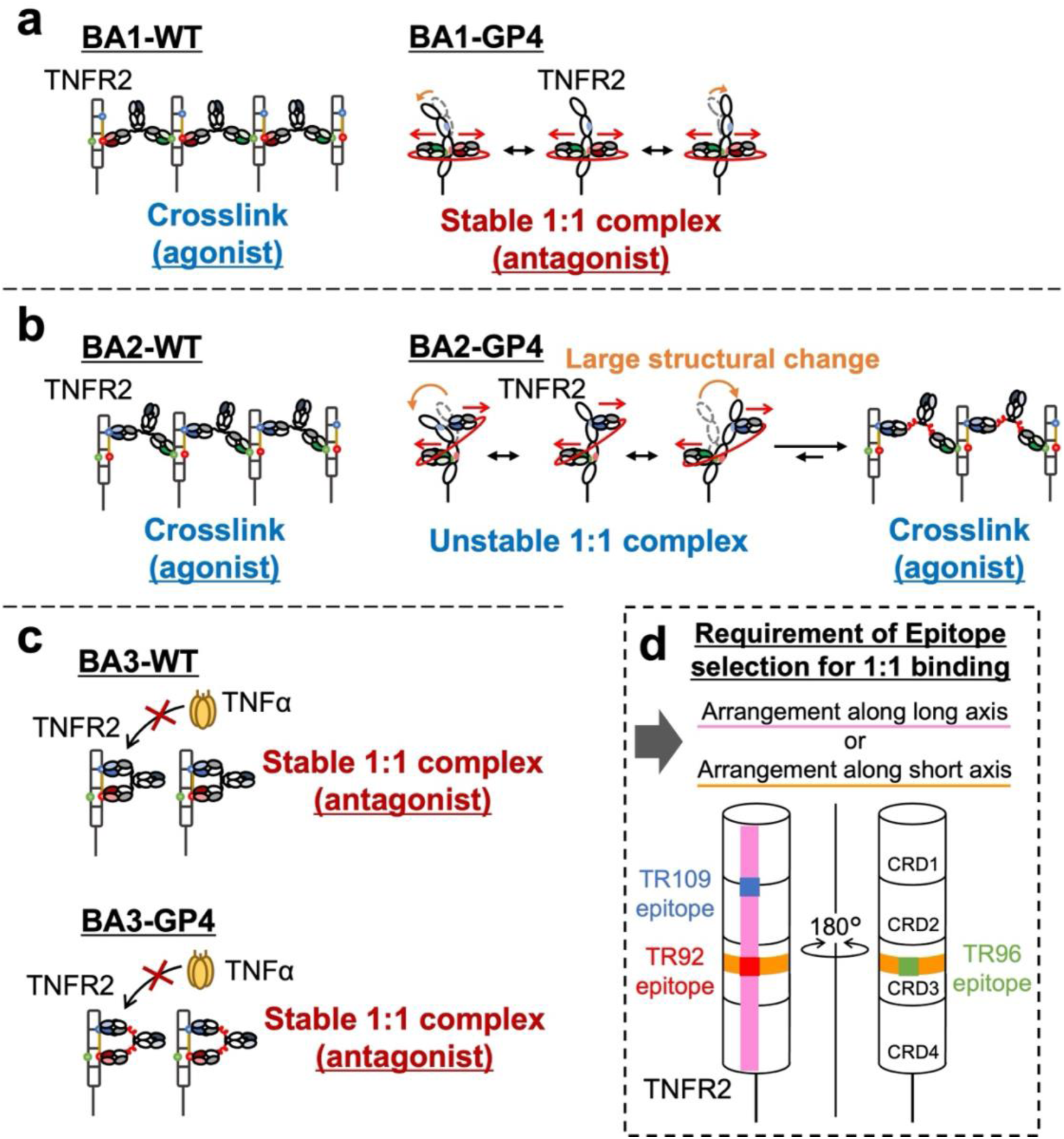
Suggested mechanism for differences in the effect of linker insertion to biparatopic antibodies (BpAbs) and implication for BpAb design. **a-c** Changes in the biological functions by inserting the peptide linker into the hinge region of BA1 (**a**), BA2 (**b**) and BA3 (**c**). **a** Linker insertion to agonistic BA1-WT recognizing the epitopes in the opposite side of TNFR2 resulted in conversion into an antagonist due to stable 1:1 complex formation. **b** Linker insertion to agonistic BA2-WT recognizing distant epitopes of TNFR2 resulted in unstable 1:1 complex formation and agonistic activity remained. **c** Linker insertion to antagonistic BA3-WT recognizing adjacent epitopes along the long axis of TNFR2 did not affect the function and 1:1 complex was maintained. **d** Selection of epitope pair required for 1:1 binding. In addition to the previously reported combination in the same side along the long axis (pink), the combination in the opposite side vertical to the axis (orange) is also allowed when a sufficiently long linker is inserted.

Understanding the mechanism by which activity conversion is achieved helps in the rational design of BpAbs to control their functions. In particular, less-effective linkers for 1:1 complex formation were informative for understanding the mechanism. The tertiary structure of the 1:1 BA1-GP4–TNFR2 complex was successfully obtained by cryo-EM single-particle analysis. The distance between the C-termini of the two Fabs is ca. 15 nm, which needs to be connected by long linkers. It was reasonable to observe little effect by introduction of short peptide linkers that cannot reach the 15 nm distance (10–32 amino acids; **Table 1** and **Supplementary Figure 7**). The length of a single unit of a GP linker, when fully stretched, is approximately 11 nm. However, due to its high flexibility, stretched conformations with restricted degrees of freedom are not dominant. Consequently, in the case of BA1-GP2 (fully stretched linker length: 22 nm), the linker length was insufficient, and the 1:1 complex would be energetically unfavorable. This caused the equilibrium to shift toward the formation of 2:2 or larger complexes (**Figure 3**), and the maximum signal-inducing activity of BA1-GP2 was reduced only by 50% of that of BA1-WT. In our previous study using synthetic linkers, we observed a similar trend where stretch close to the maximum length is insufficient to form a stable 1:1 complex^48^. Extension of linkers into BA1-GP4 (fully stretched linker length: 44 nm) and into BA1-GP8 (fully stretched linker length: 88 nm) was thus advantageous.

In the case of BA2, 50% of the maximum signal-inducing activity remained even when linkers of sufficient length were introduced (GP4 and GP8; **Figure 2b**). Immunocomplexes of 2:2 or larger were observed irrespective of the inserted linkers (**Figure 3**). Difference of BA2 from BA1 was observed in terms of the structural motions of TNFR2 in CGMD of 1:1 immunocomplexes. For BA2-GP4, the bound Fabs C-terminally linked by the linker moved in a push-and-pull motion (**Figure 5c**). This could be attributed to the epitopes of 96-Fab and 109-Fab in BA2-GP4 present in different positions along the long axis of TNFR2. In experimental conditions, where dissociation and rebinding of the Fabs to TNFR2 was allowed, such motion destabilized 1:1 complex, and alternatively, crosslinked complexes were formed (**Figure 6b**). The observation was in contrast to the complex with BA1-GP4, where 96-Fab and 92-Fab were bound at almost the same positions along the long axis of TNFR2 (**Figure 4a**), and minimal push-and-pull motion was observed (**Figure 5c**). Relatively small mobility in the case of BA1-GP4–TNFR2 complex was consistent with the formation of the stable 1:1 complex (**Figure 6a**). The difference may explain the epitope dependency observed between BA1-GP4 and BA2-GP4 in complex formation with TNFR2.

Insertion of the peptide linkers into the hinge region of BA3-WT, an antagonistic BpAb, did not affect the size of the immunocomplex or its signal-blocking activities (**Figure 6c**). This feature was consistent among BpAbs; it never happened that linker insertion resulted in stronger signal-inducing activity. Bivalent BpAbs would essentially favor the formation of smaller complexes unless any structural restriction is present.

Previously, we reported that binding to two epitopes on the same side along the long axis of TNFR2 was required to induce a 1:1 complex by IgG-like BpAbs (**Figure 6d**)^18^. The native IgG-like structure may impose restrictions on narrow epitope selection. This study provides a novel approach for addressing this structural constraint. Introduction of a long linker induces the formation of smaller immunocomplexes, even when the use of an IgG-like structure leads to cluster formation. For BA1-GP4 and BA1-GP8, the Fabs that recognized 180° opposite epitopes were linked by a linker of sufficient length. A 1:1 complex was stably formed between BA1-GP4 and TNFR2 without significantly distorting the structure of CRD1 in TNFR2. Similarly, for BpAb binding to nearly the same position along the long axis and on different sides (**Figure 6d**), a 1:1 complex can be induced by the use of a linker of sufficient length. However, when the vertical position was different, the 1:1 complex induction was only partial. The relative positions of the epitopes are crucial, even for linker-extended BpAbs. This could contribute to the development of an efficient design strategy of 1:1-binding BpAbs against membrane proteins.

In conclusion, agonistic BpAbs were converted into antagonists via linker insertion. Both the combination of epitopes and the length of the linker are important for inducing the formation of the 1:1 complex that confers this activity. The BpAb design strategy based on this study can be widely applied to other single-transmembrane receptors, including TNFRSF. The findings of the present study on the molecular recognition mechanisms could also contribute to the design of a wide range of biotherapeutics, such as peptides, alternative scaffold proteins, or nucleic acid aptamers, and contribute to expanding the therapeutic potential of BpAbs and these modalities.

## Materials and methods

### Cloning, expression, and purification of recombinant antibodies

Heavy and light chains of BpAbs were designed in the CrossMAb format^38^ with the amino-acid sequences shown in **Supplementary Tables 1-3**. The DNA encoding the heavy and light chains that constitute BpAbs were cloned into pFUSE-CHIg-hG1 and pFUSE2-CLIg-hκ (InvivoGen), respectively. BpAbs were produced by the transfection of four plasmids in equal masses using the ExpiCHO Expression System (Thermo Fisher Scientific, Waltham, MA, USA), according to the manufacturer’s standard protocol. The cells were removed by centrifugation at 6,000 × *g* for 15 min and the culture supernatant was filtered through a 0.20 µm filter. The BpAbs were captured on an rProtein A Sepharose Fast Flow column (Cytiva). The column was washed with 10 mL of phosphate-buffered saline PBS (pH 7.4) and eluted with 100 mM sodium citrate pH 3.0 (3.0 mL). The eluent was dialyzed in PBS. BpAbs without linker insertion or with one linker (upper half of Table 1) were purified through size-exclusion chromatography (SEC) using a HiLoad 16/600 Superdex 200 pg column (Cytiva) in PBS. For BpAbs with two linkers inserted into both heavy chains (lower half of Table 1), SEC was conducted using Superdex 200 Increase 10/300 GL column. Then, cation-exchange chromatography (CIEX) was conducted using a Mono S 5/50 GL column (Cytiva) with Buffer A (20 mM MES-NaOH, 10 mM NaCl, pH 6.0) and Buffer B (20 mM MES-NaOH, 1 mM NaCl, pH 6.0). The BpAbs were eluted using a linear gradient from 0%B to 50%B over 50 min. The BpAbs were dialyzed and stored in PBS.

Recombinant Fab proteins with a C-terminal hexahistidine tag were expressed and purified as previously reported^48^. Briefly, proteins were expressed using the Expi293 Expression System (Themo Fisher Scientific), and the culture supernatant was dialyzed with 20 mM Tris-HCl and 300 mM NaCl (Buffer C). Fab proteins were captured in 1 mL of cOmplete His-Tag Purification Resin (Roche Diagnostics). The column was washed with Buffer C containing 5, 10, or 20 mM imidazole (2.5 mL each) and eluted with Buffer C containing 200 or 500 mM imidazole (2.5 mL each). The eluent was dialyzed against buffer C and subjected to SEC using a HiLoad Superdex200 16/600 column (Cytiva) in the same buffer. The fractions containing monomeric Fab proteins were purified using CIEX following the protocol described above.

### Deglycosylation of GP linker inserted into BpAbs

BpAb (1 mg), dialyzed to PBS after Protein A affinity chromatography, was mixed with 2 μL *O*-glycosidase (#P0733S, New England BioLabs), 2 μL α2-3,6,8 neuraminidase (#P0720S, New England BioLabs), 15 μL of 10× GlycoBuffer 2 (provided with *O*-glycosidase) and deionized-distilled water to make a 150 μL reaction volume. After 4 h incubation at 37 ℃, Protein A affinity chromatography was conducted to remove the enzymes, and the eluent was further purified by SEC and CIEX following the same protocol for the original BpAbs.

### Reporter gene assay

Human TNFR2-expressing Ramos-Blue cells were cultured in IMDM supplemented with 2 mM glutamine and 10% fetal bovine serum (FBS) as previously described^18^. Cells were seeded at 1 × 10^5^ cells in 50 μL of medium in the absence or presence of 50 ng/mL TNFα (#AF-300-01A, Peprotech USA). Antibodies at the indicated concentrations (1.2 pM–400 nM in a 3.16-fold dilution series) were added to the cells. The cells were incubated for 18 h and the secreted alkaline phosphatase was analyzed using *p*-nitrophenyl phosphate (1.7 mg/mL). Colorimetric changes were determined by measuring absorbance at 405 nm using an EnVision microplate reader (PerkinElmer, Waltham, MA, USA). Signals were normalized to the average absorbance of the six wells incubated without antibody or TNFα (negative) and the six wells incubated only with 50 ng/mL TNFα (positive). For the chart showing the NF-κB signals, the negative and positive values were set at 0.1 and 1.0, respectively.

### Flow cytometry

TNFR2-expressing Ramos-Blue cells (5 × 10^4^ cells/condition) were resuspended with 50 µL of 3.16-fold serial dilution series of BpAbs in FCM buffer (PBS containing 0.2% sodium azide and 5% FBS) and incubated on ice for 1 h. After washing twice, the cells were resuspended with 25 µL of PE-goat anti-human IgG (1/400 dilution, #109-116-170, Jackson ImmunoReseach, USA) in FCM buffer and incubated on ice for 30 min. The cells were washed once, resuspended in FCM buffer, and analyzed using a CytoFLEX (Beckman Coulter Life science, USA). Data were analyzed using CyExpert (Beckman Coulter Life science, USA).

### Size determination of immunocomplexes

TNFR2-MBP and sTNFR2 were obtained as previously reported^18^. SEC of the immunocomplexes was performed in PBS using a Superose 6 Increase 10/300 GL column (Cytiva) run at 0.45 mL/min. The mixture of BpAb (1 μM) and TNFR2-MBP (1 μM) in PBS, 100 μL, was analyzed. Mass photometry was conducted using Refeyn One (Refeyn Ltd., Oxford, UK). Five microliter solution containing BpAb (111 nM) and sTNFR2 (22 nM) was diluted four-fold with 15 μL PBS pre-loaded for hydration on a glass slide. Interferometric scattering was observed under a microscope for 1–2 min, and the accumulated interference signal was analyzed using the software provided by the manufacturer in the same manner as the literature^49^.

### Sample preparation for cryo-electron microscopy

The 1:1 complex of BA1-GP4 and TNFR2-MBP was purified from the mixture of BA1-GP4 (17.8 μM, 70 μL) and TNFR2-MBP (27.9 μM, 56 μL) using Superose 6 Increase 10/300 GL column. Fractions corresponding to 1:1 complex were collected and concentrated by ultrafiltration to 0.71 mg/mL (Amicon-Ultra-4, 10 K). Three microliters of the complex solution (0.35 mg/mL) were applied to a Quantifoil R1.2/1.3 Cu 200 mesh grid (Quantifoil Micro Tools GmbH, Jena, Germany), which was glow-discharged for 20 s at 20 mA using a JEC-3000FC sputter coater (JEOL). The grid was blotted with a blot force of –10 and a blot time of 2 s in a Vitrobot Mark IV chamber (Thermo Fisher Scientific) equilibrated at 4 °C and 100% humidity, and then immediately plunged into liquid ethane. Excess ethane was removed using filter paper, and the grids were stored in liquid nitrogen. The image dataset was collected using a SerialEM^50^, yoneoLocr^51^, and JEM-Z300FSC (CRYO ARM 300: JEOL) operated at 300 kV with a K3 direct electron detector (Gatan, Pleasanton, CA, USA) in the CDS mode. A Ω-type in-column energy filter was operated with a slit width of 20 eV for zero-loss imaging. The nominal magnification was 60,000×, corresponding to an image pixel size of 0.88 Å. The defocus varied between −0.5 and −2.0 µm. Each movie was fractionated into 40 frames (0.081 s each, total exposure: 4.87 s), with a total dose of 80 e^−^/Å^2^.

### Cryo-EM image processing and model building

A gain reference image was prepared with the relion_estimate_gain command in RELION 4.0^52^ using the first 500 movies. The images were processed using cryoSPARC ver. 4.2.1^53^. A total of 5,508 movies were imported, and motion-corrected and contrast transfer functions (CTFs) were estimated. A total of 5,172 micrographs with maximum CTF resolutions greater than 5 Å were selected. First, the particles were automatically picked using a blob picker job with a particle diameter between 100 and 200 Å. After particle extraction with 2x binning, a 2D classification into 100 classes was performed to select clear 2D class averages as templates for subsequent particle picking. Then, the particles were again automatically picked with the templates, and 2,971,529 particle images were extracted with a box size of 360 pixels using 2x binning (downscaled to 180 pixels). Two rounds of 2D classification into 100 classes were conducted to select 409,348 particles. The number of final full iterations and the batch size per class were increased to 20 and 400, respectively. The selected particles were extracted again with a box size of 360 pixels into 300 pixels (1.048 Å/pix), and subjected to Ab-Initio Reconstruction. The best initial model was selected from the two reconstructed models, and the handedness of the map and mask was flipped. After non-uniform refinement, global/local CTF refinement and non-uniform refinement^54^ were performed, a final map was reconstructed at 3.73 Å resolution (FSC = 0.143). Homology models of 109-Fab and 92-Fab were generated using the SWISS-MODEL^55^. The atomic model of TNFR2 (PDB entry: 3ALQ) and the homology models of the Fabs were manually fitted into the density and modified using UCSF Chimera^56^ and Coot^57^. Realspace refinement was performed using the PHENIX software^58^.

### Analysis of amino-acid residues of TNFR2 in the interface with 92-Fab from two different BpAbs

The amino acid residues at the binding interface between TNFR2 and 92-Fab in BA1-GP4 (PDB ID: 9LFL) or Bp109-92 (PDB ID: 8HLB) were analyzed using the PISA server^46^. The change in buried surface area by interaction (ΔBSA) in the interface is calculated. Interface residues are defined as ΔBSA > 0.

### Modeling initial structure of immunocomplexes for coarse-grained molecular dynamics (CGMD) simulation

The initial structures for CGMD were modeled using Molecular Operating Environment (MOE) version 2022.2 (Chemical Computing Group Inc., Canada) software. In the case of BA1-GP2 and BA1-GP4, initial structure of tertiary complex of TNFR2, TR92-Fab and TR96-Fab was adopted from PDB ID 9LFL. The linkers were introduced following the method shown in **Supplementary Figures 24 and 25**. In the case of BA2-GP2 and BA2-GP4, PDB entries 8HLB and 9LFL were superimposed by TNFR2, and duplicated amino acid residues were removed. Arg119 in 8HLB and Lys200 in 9LFL were then joined to create the initial tertiary structure of TNFR2, TR109-Fab and TR96-Fab (**Supplementary Figure 26**). The linkers were introduced following the method shown in **Supplementary Figures 27 and 28**. In all cases, the structure of Fc was modeled in MOE.

### Coarse-Grained Molecular Dynamics (CGMD) simulations

As course-grained (CG) model, we adopted the atomic interaction-based CG (AICG2+) model^59^, in which each amino acid in proteins is represented by a CG particle located at the position of Cα atom. The group of parameters related to AICG2+ basically adapts the default parameters of CafeMol ver 3.2.1 (https://www.cafemol.org). The CGMD simulation was conducted at *T* = 310.15 K for 10^6^ steps with a time step *dt* = 0.1 for each parameter set. All disulfide bonds in the structure were bound together by harmonic springs. The strength of the bonding is *k_ij_(r_ij_-l_o_)*^2^. *k_ij_*, *r_ij_*and *l_o_* represent the spring constant (kcal/mol/Å), the distance of two mass points (Å) and the bond length (Å). The value of *k_ij_*was 20.0 kcal/mol/Å^2^ and the value of *l_o_* was 5 Å. Ten runs of the CGMD simulations were conducted using CafeMol for four different sample trajectories. The coordinates of Cα atoms of TNFR2 were extracted from the runs, and were fitted to a reference structure.

### Principal component analysis (PCA) of TNFR2 structure

PCA was performed to analyze structural changes in TNFR2 using GROMACS version 2023.3^60^. The covariance matrix for the fitted data is then computed. PCA was performed on the fitted data and eigenvectors were computed.

## Acknowledgment

We thank Dr. Satoshi Nagata for the reporter cells, Dr. Takao Arimori and Dr. Kohei Shiba for assistance with mass photometry, Dr. Shinsuke Inuki for assistance with flow cytometry, Mr. Toshiki Fujii for assistance with CGMD calculations, Dr. Miki Kinoshita for assistance in structure refinement, Dr. Takayuki Kato for assistance in database deposition, and Ms. Reiko Satoh for assistance in plasmid vector preparation. This work was partially supported by AMED grant numbers JP22ak0101099 (H.K. and H.A.), JP24ama121042 (H.K. and H.A.), JP23ama121003 (K.N.), JSPS grant number JP24K01270 (H.A.), JEOL YOKOGUSHI Research Alliance Laboratories of Osaka University (K.N.), the Takeda Science Foundation (H.A.), and the Kyoto University Foundation (H.A.). T.O. was supported by the Graduate Program for Medical Innovation, Kyoto University.

## Conflict of Interest Statement

H.A. is a scientific advisor at Epitope Science Co., Ltd.. H.K. is a board member and shareholder of Epitope Science Co., Ltd.. The other authors declare no conflicts of interest.

## Author Contribution

H.A. conceived of the study. T.O., H.O., and H.A. designed the study. T.O. and J.F. conducted the experiments. S.M., T.M., and R.K. analyzed the data. K.N., R.K., and Y.O. developed the methodology. T.O. and H.A. wrote the original manuscript. S.M., R.K., H.K., and H.O. revised the manuscript. All the authors agree to submit this manuscript for publication.

## Notes

### Summary of Updates

Items in Figure 5b were wrongly labeled and they were revised.

