## Supplementary Information for "Conversion of an agonistic anti-TNFR2 biparatopic antibody into an antagonist by insertion of peptide linkers into the hinge region"

**Table of Contents**


| Contents | Title | Page # |
|---|---|---|
| Supplementary Table 1 | Amino acid sequence of BA1 | S4 |
| Supplementary Table 2 | Amino acid sequence of BA2 | S5 |
| Supplementary Table 3 | Amino acid sequence of BA3 | S6 |
| Supplementary Table 4 | Cryo-EM data collection and processing | S7 |
| Supplementary Table 5 | Eigenvalues and contribution of 1st-5th Principal Components (PC1-5) | S7 |
| Supplementary Figure 1 | Format exchange from intein-mediated protein trans-splicing (IMPTS) into CrossMAb | S8 |
| Supplementary Figure 2 | Preparative size-exclusion chromatograms of BA1 inserted with a peptide linker into one of heavy chains | S9 |
| Supplementary Figure 3 | Preparative size-exclusion chromatograms of BA2 inserted with a peptide linker into one of heavy chains | S10 |
| Supplementary Figure 4 | Preparative size-exclusion chromatograms of BA3 inserted with a peptide linker into one of heavy chains | S11 |
| Supplementary Figure 5 | SDS-PAGE analysis of BpAbs | S12 |
| Supplementary Figure 6 | Decomposition of proteins stored at 4 °C, one month after purification | S13 |
| Supplementary Figure 7 | Control of biological activities through linker insertion into the hinge region of one of heavy chains | S14 |
| Supplementary Figure 8 | Preparative size-exclusion chromatograms of BA1-WT, BA1-GP2, BA1-GP4 and BA1-GP8 | S15 |
| Supplementary Figure 9 | Preparative cation-exchange chromatograms of BA1-WT, BA1-GP2, BA1-GP4 and BA1-GP8 | S16 |
| Supplementary Figure 10 | Preparative size-exclusion chromatograms of BA2-WT, BA2-GP2, BA2-GP4 and BA2-GP8 | S17 |
| Supplementary Figure 11 | Preparative cation-exchange chromatograms of BA2-WT, BA2-GP2, BA2-GP4 and BA2-GP8 | S18 |
| Supplementary Figure 12 | Agonistic and antagonistic activities of the mixtures of three combinations of two Fab in a reporter gene assay | S19 |
| Supplementary Figure 13 | Preparative size-exclusion chromatograms of BA3-WT, BA3-GP2 and BA3-GP4 | S20 |
| Supplementary Figure 14 | Preparative cation-exchange chromatograms of BA2-GP2 and BA2-GP4 | S21 |

| | | |
|---|---|---|
| Supplementary Figure 15 | Agonistic and antagonistic activities of BA3-WT, BA3 GP2 and BA3-GP4 in a reporter gene assay | S21 |
| Supplementary Figure 16 | Preparative size-exclusion and cation-exchange chromatograms of BA1-GP2, BA1-GP4, BA2-GP2 and BA2-GP4 after deglycosylation | S22 |
| Supplementary Figure 17 | Agonistic and antagonistic activities of BpAbs after deglycosylation in a reporter gene assay. | S23 |
| Supplementary Figure 18 | Apparent binding affinity of BpAbs to TNFR2-expressing Ramos-Blue cells analyzed by using flow cytometry | S23 |
| Supplementary Figure 19 | Immunocomplex formed between BA3 and TNFR2 | S24 |
| Supplementary Figure 20 | Sample preparation for cryo-electron microscopy | S24 |
| Supplementary Figure 21 | Image processing of BA1-GP4/TNFR2-MBP complex | S25 |
| Supplementary Figure 22 | Amino-acid residues of TNFR2 in the interface with 92-Fab from two different BpAbs | S26 |
| Supplementary Figure 23 | The epitope of 96-Fab binding to TNFR2 | S27 |
| Supplementary Figure 24 | Strategy for building the initial structure of the complexes of BA1-GP2 and TNFR2 | S28 |
| Supplementary Figure 25 | Strategy for building the initial structure of the complexes of BA1-GP4 and TNFR2 | S29 |
| Supplementary Figure 26 | Strategy for building ternary complex of 109-Fab, 96-Fab and TNFR2 | S30 |
| Supplementary Figure 27 | Strategy for building the initial structure of the complexes of BA2-GP2 and TNFR2 | S31 |
| Supplementary Figure 28 | Strategy for building the initial structure of the complexes of BA2-GP4 and TNFR2 | S32 |

**Supplementary Table 1.** Amino acid sequence of BA1[a]

| Knob chain | Hole chain |
|---|---|
| ***HC*** | ***HC*** |
| KVQLQQSGAELVKPGASVKLSCKASGYTFTESIIHWVKQRSGQGLEWIGW<br>FYPGSDNINYNEKFKDKATLTADKSSSTVYMELTRLTSEDSAVYFCASHE<br>GPYVYFDYWGQGTTLTVSSASTKGPSVFPLAPSSKSTSGGTAALGCLVKD<br>YFPEPVTVSWNSGALTSGVHTFPAVLQSSGLYSLSSVVTVPSSSLGTQTY<br>ICNVNHKPSNTKVDKKVEPKS**---Hinge region---**PAPELLGGPSV<br>FLFPPKPKDTLMISRTPEVTCVVVDVSHEDPEVKFNWYVDGVEVHNAKTK<br>PREEQYNSTYRVVSVLTVLHQDWLNGKEYKCKVSNKALPAPIEKTISKAK<br>GQPREPQVYTLPPCRDELTKNQVSLWCLVKGFYPSDIAVEWESNGQPENN<br>YKTTPPVLDSDGSFFLYSKLTVDKSRWQQGNVFSCSVMHEALHNHYTQKS<br>LSLSPGK | EVQLQQSGAELVKPGASVKLSCTPSGFNIKDTYMHWVKQRPEQGLEWIGR<br>IDPANGYTEYDPKFQDKATITADTSSNTAYLQLSSLTSEDTAVYYCADTQ<br>LYYWGQGTTLTVSSASVAAPSVFIFPPSDEQLKSGTASVVCLLNNFYPRE<br>AKVQWKVDNALQSGNSQESVTEQDSKDSTYSLSSTLTLSKADYEKHKVYA<br>CEVTHQGLSSPVTKSFNRGE**---Hinge region---**PAPELLGGPSVF<br>LFPPKPKDTLMISRTPEVTCVVVDVSHEDPEVKFNWYVDGVEVHNAKTKP<br>REEQYNSTYRVVSVLTVLHQDWLNGKEYKCKVSNKALPAPIEKTISKAKG<br>QPREPQVCTLPPSRDELTKNQVSLSCAVKGFYPSDIAVEWESNGQPENNY<br>KTTPPVLDSDGSFFLVSKLTVDKSRWQQGNVFSCSVMHEALHNHYTQKSL<br>SLSPGK |
| ***LC*** | ***LC*** |
| DIVMTQSHKFMSTSVGDRVSITCKASQDVSTAVAWYQQKPGQSPKLLIYW<br>TSTRHTGVPDRFTGSGSGTDYTLTISSVQAEDLALYYCQHHYSTPYTFGG<br>GTKLEIQRTVAAPSVFIFPPSDEQLKSGTASVVCLLNNFYPREAKVQWKV<br>DNALQSGNSQESVTEQDSKDSTYSLSSTLTLSKADYEKHKVYACEVTHQG<br>LSSPVTKSFNRGEC | QIVLTQSPAIMSASLGERVTMTCTASSSVSSTYLHWYQQKPGSSPKLWIY<br>STSNLASGVPARFSGSGSGTSYSLTISNMEAEDAATYYCHQYHRSPLTFG<br>AGTKLELKSSASTKGPSVFPLAPSSKSTSGGTAALGCLVKDYFPEPVTVS<br>WNSGALTSGVHTFPAVLQSSGLYSLSSVVTVPSSSLGTQTYICNVNHKPS<br>NTKVDKKVEPKSC |

[a] Disulfide-linked knob-into-hole mutations are highlighted in cyan. See Table 1 for the hinge region sequences.

**Supplementary Table 2.** Amino acid sequence of BA2[a]

| Knob chain | Hole chain |
|---|---|
| ***HC***<br>EVQLQQSGAELVKPGASVKLSCTPSGFNIKDTYMHWVKQRPEQGLEWIGR<br>IDPANGYTEYDPKFQDKATITADTSSNTAYLQLSSLTSEDTAVYYCADTQ<br>LYYWGQGTTLTVSSASTKGPSVFPLAPSSKSTSGGTAALGCLVKDYFPEP<br>VTVSWNSGALTSGVHTFPAVLQSSGLYSLSSVVTVPSSSLGTQTYICNVN<br>HKPSNTKVDKKVEPKS**---Hinge Region---**PAPELLGGPSVFLFPP<br>KPKDTLMISRTPEVTCVVVDVSHEDPEVKFNWYVDGVEVHNAKTKPREEQ<br>YNSTYRVVSVLTVLHQDWLNGKEYKCKVSNKALPAPIEKTISKAKGQPRE<br>PQVYTLPPCRDELTKNQVSLWCLVKGFYPSDIAVEWESNGQPENNYKTTP<br>PVLDSDGSFFLYSKLTVDKSRWQQGNVFSCSVMHEALHNHYTQKSLSLSP<br>GK | ***HC***<br>EQVQLKESGPGLVAPSQSLSITCTVSGFSLTVYGVNWVRQPPGKGLEWLG<br>MIWGDGSTAYNSALKSRLTITKDNSKTQVFLKMNSLQTDDTARYYCARDG<br>RRYALDYWGQGTSVTVSSASVAAPSVFIFPPSDEQLKSGTASVVCLLNNF<br>YPREAKVQWKVDNALQSGNSQESVTEQDSKDSTYSLSSTLTLSKADYEKH<br>KVYACEVTHQGLSSPVTKSFNRGE**---Hinge region---**PAPELLGG<br>PSVFLFPPKPKDTLMISRTPEVTCVVVDVSHEDPEVKFNWYVDGVEVHNA<br>KTKPREEQYNSTYRVVSVLTVLHQDWLNGKEYKCKVSNKALPAPIEKTIS<br>KAKGQPREPQVCTLPPSRDELTKNQVSLSCAVKGFYPSDIAVEWESNGQP<br>ENNYKTTPPVLDSDGSFFLVSKLTVDKSRWQQGNVFSCSVMHEALHNHYT<br>QKSLSLSPGK |
| ***LC***<br>QIVLTQSPAIMSASLGERVTMTCTASSSVSSTYLHWYQQKPGSSPKLWIY<br>STSNLASGVPARFSGSGSGTSYSLTISNMEAEDAATYYCHQYHRSPLTFG<br>AGTKLELKRTVAAPSVFIFPPSDEQLKSGTASVVCLLNNFYPREAKVQWK<br>VDNALQSGNSQESVTEQDSKDSTYSLSSTLTLSKADYEKHKVYACEVTHQ<br>GLSSPVTKSFNRGEC | ***LC***<br>DIVLTQSPTSLAVSLGQRATISCRASESVDSYGDSFLHWYQQKPGQPPIL<br>LIYRASNLDSGIPARFSGSGSRTDFTLTINPVEADDVATYYCQQSNEDPY<br>TFGGGTKVTVLSSASTKGPSVFPLAPSSKSTSGGTAALGCLVKDYFPEPV<br>TVSWNSGALTSGVHTFPAVLQSSGLYSLSSVVTVPSSSLGTQTYICNVNH<br>KPSNTKVDKKVEPKSC |

[a] Disulfide-linked knob-into-hole mutations are highlighted in cyan. See Table 1 for the hinge region sequences.

**Supplementary Table 3.** Amino acid sequence of BA3[a]

| Knob chain | Hole chain |
| --- | --- |
| ***HC***<br>KVQLQQSGAELVKPGASVKLSCKASGYTFTESIIHWVKQRSGQGLEWIGW<br>FYPGSDNINYNEKFKDKATLTADKSSSTVYMELTRLTSEDSAVYFCASHE<br>GPYVYFDYWGQGTTLTVSSASTKGPSVFPLAPSSKSTSGGTAALGCLVKD<br>YFPEPVTVSWNSGALTSGVHTFPAVLQSSGLYSLSSVVTVPSSSLGTQTY<br>ICNVNHKPSNTKVDKKVEPKS**---Hinge region---**PAPELLGGPSV<br>FLFPPKPKDTLMISRTPEVTCVVVDVSHEDPEVKFNWYVDGVEVHNAKTK<br>PREEQYNSTYRVVSVLTVLHQDWLNGKEYKCKVSNKALPAPIEKTISKAK<br>GQPREPQVYTLPPCRDELTKNQVSLWCLVKGFYPSDIAVEWESNGQPENN<br>YKTTPPVLDSDGSFFLYSKLTVDKSRWQQGNVFSCSVMHEALHNHYTQKS<br>LSLSPGK | ***HC***<br>QVQLKESGPGLVAPSQSLSITCTVSGFSLTVYGVNWVRQPPGKGLEWLGM<br>IWGDGSTAYNSALKSRLTITKDNSKTQVFLKMNSLQTDDTARYYCARDGR<br>RYALDYWGQGTSVTVSSASVAAPSVFIFPPSDEQLKSGTASVVCLLNNFY<br>PREAKVQWKVDNALQSGNSQESVTEQDSKDSTYSLSSTLTLSKADYEKHK<br>VYACEVTHQGLSSPVTKSFNRGE**---Hinge region---**PAPELLGGP<br>SVFLFPPKPKDTLMISRTPEVTCVVVDVSHEDPEVKFNWYVDGVEVHNAK<br>TKPREEQYNSTYRVVSVLTVLHQDWLNGKEYKCKVSNKALPAPIEKTISK<br>AKGQPREPQVCTLPPSRDELTKNQVSLSCAVKGFYPSDIAVEWESNGQPE<br>NNYKTTPPVLDSDGSFFLVSKLTVDKSRWQQGNVFSCSVMHEALHNHYTQ<br>KSLSLSPGK |
| ***LC***<br>DIVMTQSHKFMSTSVGDRVSITCKASQDVSTAVAWYQQKPGQSPKLLIYW<br>TSTRHTGVPDRFTGSGSGTDYTLTISSVQAEDLALYYCQHHYSTPYTFGG<br>GTKLEIQRTVAAPSVFIFPPSDEQLKSGTASVVCLLNNFYPREAKVQWKV<br>DNALQSGNSQESVTEQDSKDSTYSLSSTLTLSKADYEKHKVYACEVTHQG<br>LSSPVTKSFNRGEC | ***LC***<br>DIVLTQSPTSLAVSLGQRATISCRASESVDSYGDSFLHWYQQKPGQPPIL<br>LIYRASNLDSGIPARFSGSGSRTDFTLTINPVEADDVATYYCQQSNEDPY<br>TFGGGTKVTVLSSASTKGPSVFPLAPSSKSTSGGTAALGCLVKDYFPEPV<br>TVSWNSGALTSGVHTFPAVLQSSGLYSLSSVVTVPSSSLGTQTYICNVNH<br>KPSNTKVDKKVEPKSC |

[a] Disulfide-linked knob-into-hole mutations are highlighted in cyan. See Table 1 for the hinge region sequences.

**Supplementary Table 4.** Cryo-EM data collection and processing

| Dataset | 1:1 complex of BA1-GP4 and TNFR2 |
|---|---|
| EMDB accession code | EMD-63050 |
| PDB accession code | 9LFL |
| Magnification | 60,000 |
| Voltage (kV) | 300 |
| Electron exposure ($e^{-}$/Å$^2$) | 80 |
| Defocus range (μm) | −0.5 to −2.0 |
| Pixel size (Å) | 0.878 |
| Symmetry imposed | C1 |
| Imported movies (no.) | 5,508 |
| Initial particle images (no.) | 2,971,529 |
| Final particle images (no.) | 178,242 |
| Map resolution (Å) | 3.73 |
| FSC threshold | 0.143 |

**Supplementary Table 5.** Eigenvalues and the contribution of the 1st–5th principal components (PC1-5)

| PC | value | contribution |
|---|---|---|
| 1 | 101.023 | 65.7% |
| 2 | 25.4846 | 16.6% |
| 3 | 18.4627 | 12.0% |
| 4 | 4.56841 | 3.0% |
| 5 | 1.03969 | 0.7% |

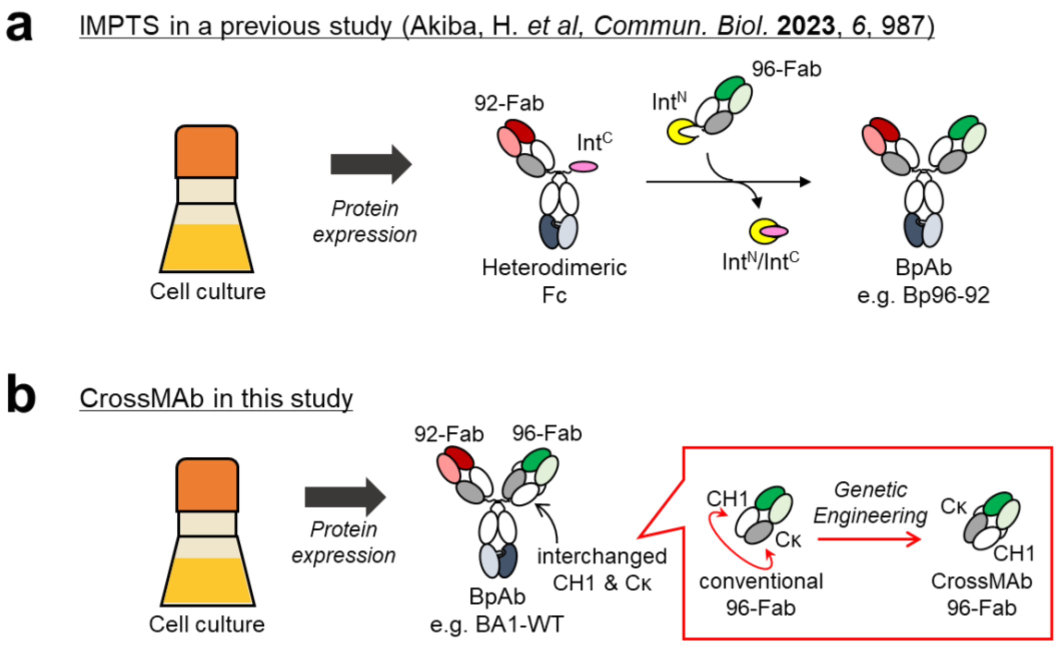


**Supplementary Figure 1.** Format transformation from intein-mediated protein trans-splicing (IMPTS) into CrossMAb.

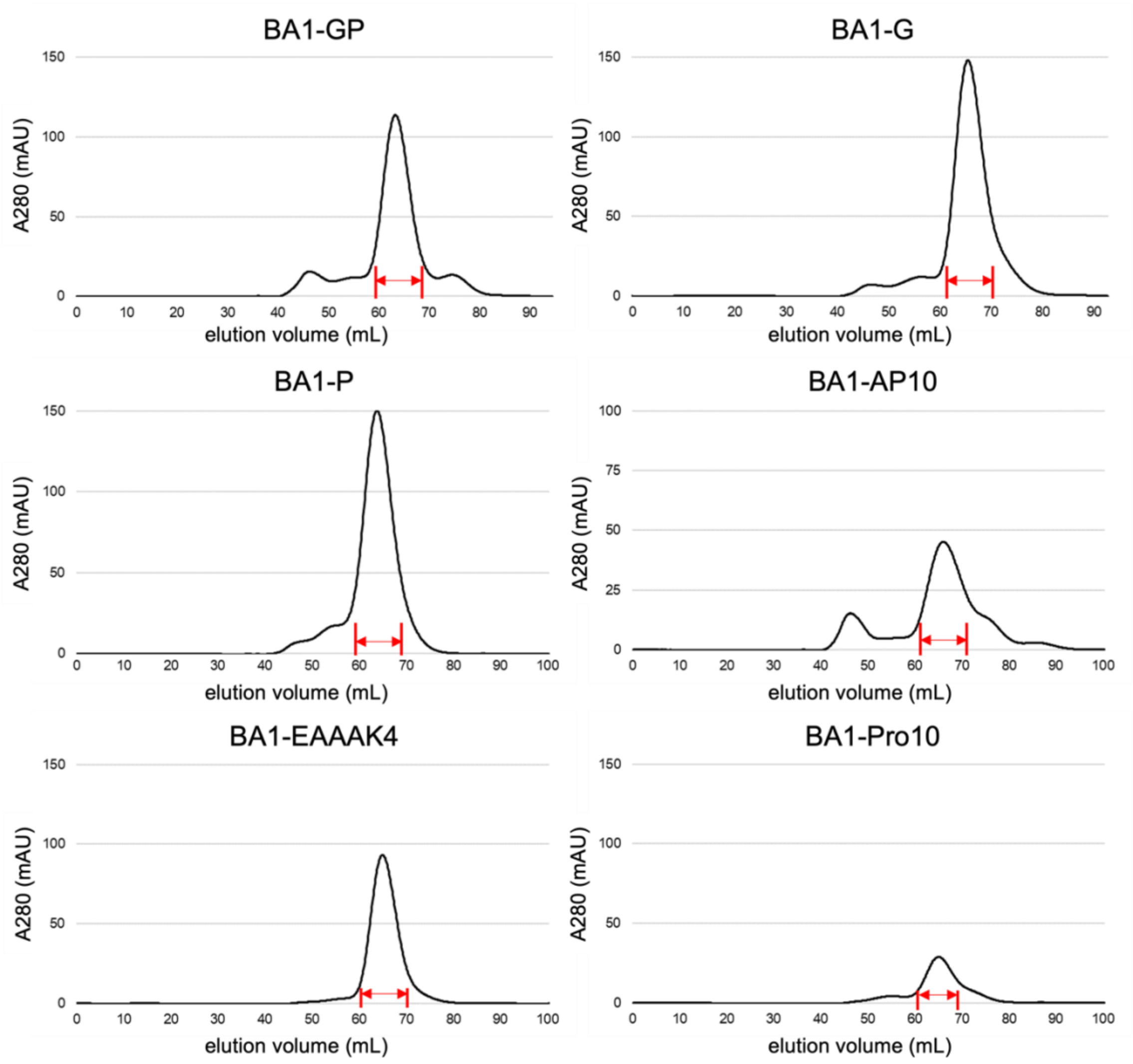


**Supplementary Figure 2.** Preparative size-exclusion chromatograms of BA1 inserted with a peptide linker into one of heavy chains. The fractions with red arrows were collected.

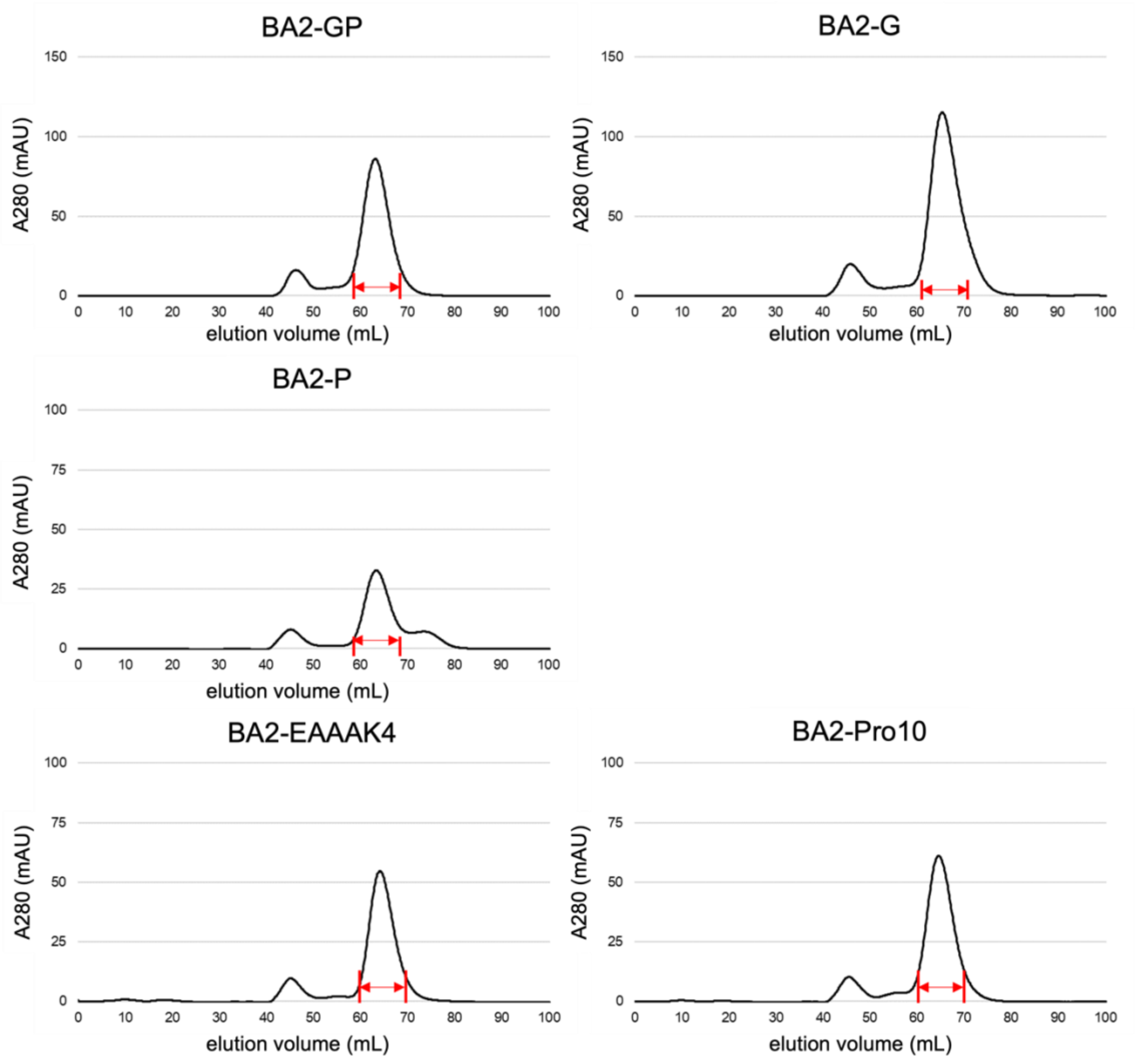


**Supplementary Figure 3.** Preparative size-exclusion chromatograms of BA2 inserted with a peptide linker into one of heavy chains. The fractions with red arrows were collected.

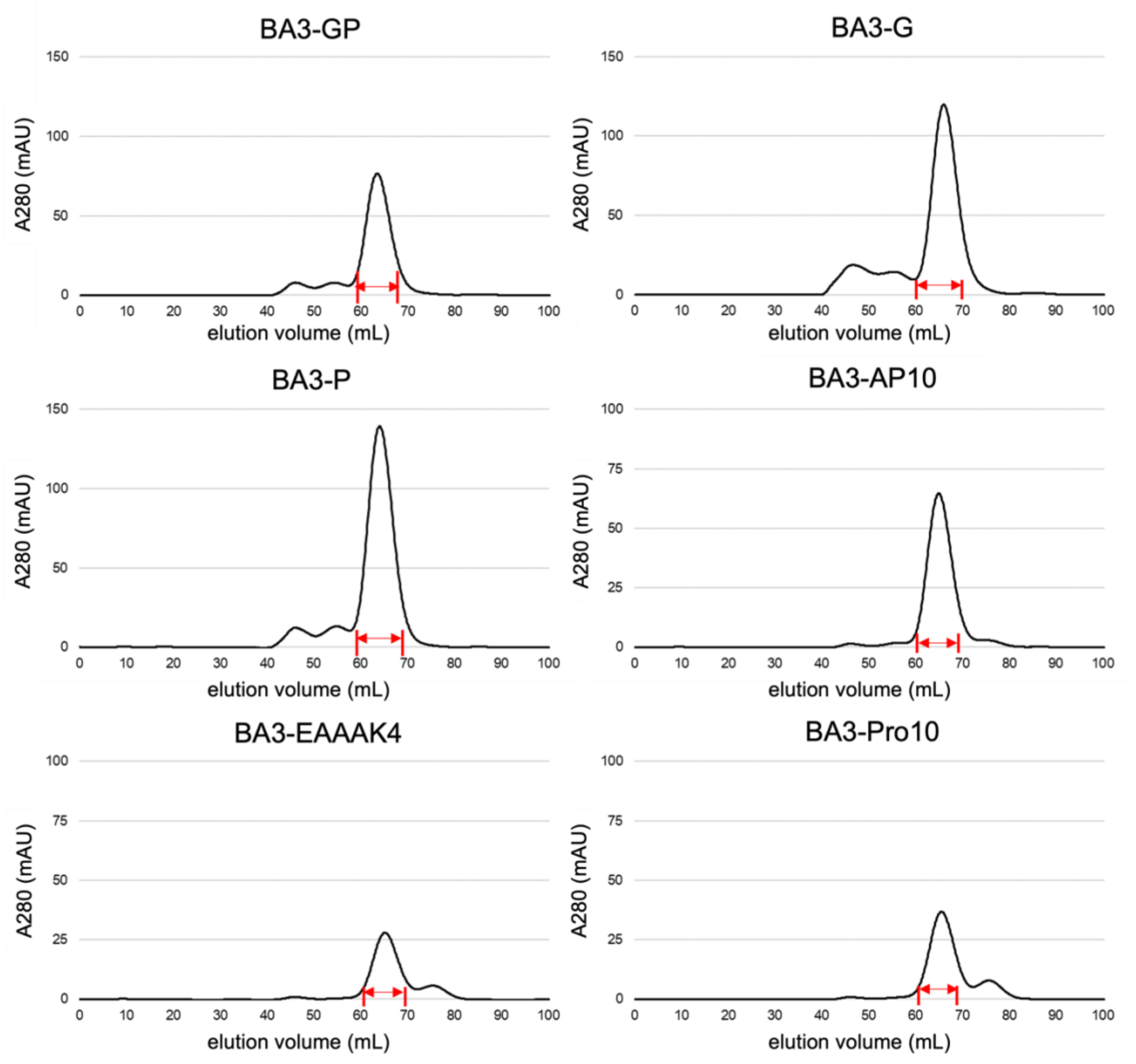


**Supplementary Figure 4.** Preparative size-exclusion chromatograms of BA3 inserted with a peptide linker into one of heavy chains. The fractions with red arrows were collected.

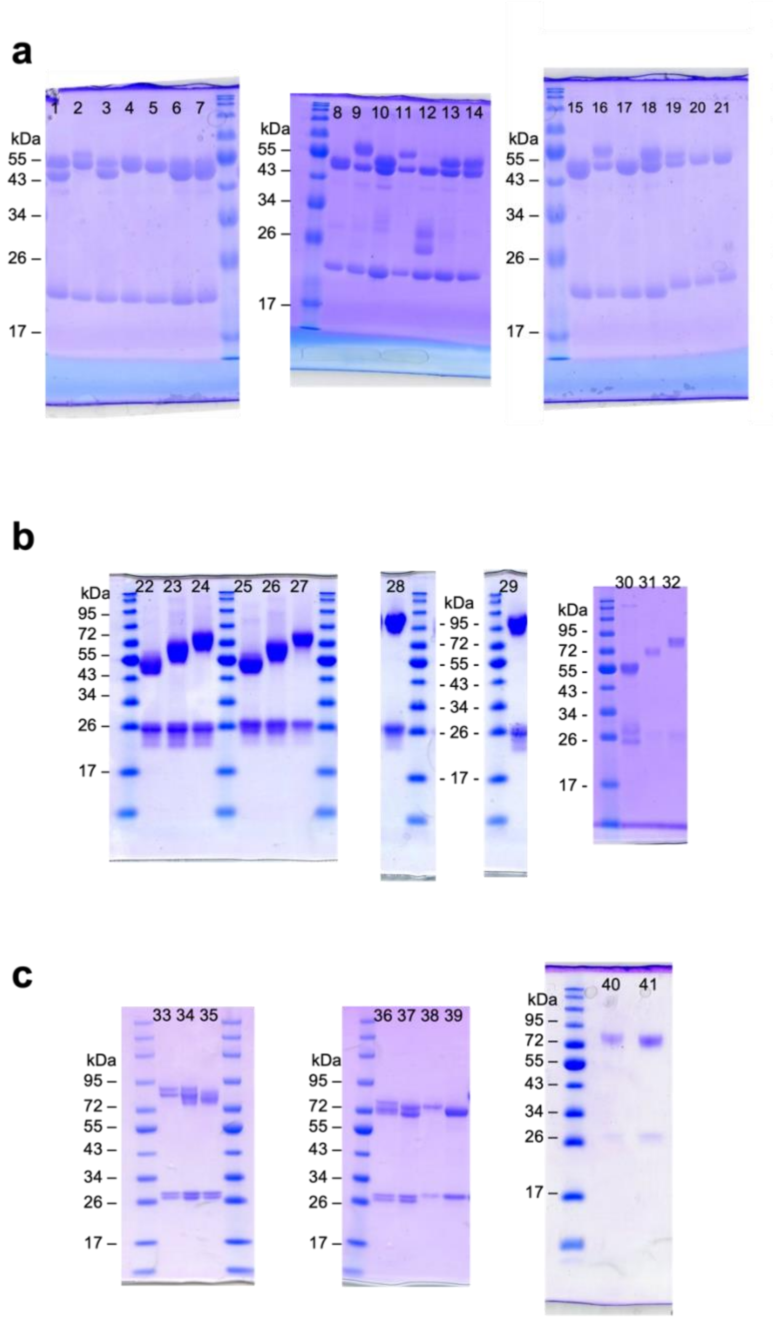


| Lane No. | Protein |
|---|---|
| 1 | BA1-WT |
| 2 | BA1-GP |
| 3 | BA1-G |
| 4 | BA1-P |
| 5 | BA1-AP10 |
| 6 | BA1-EAAAK4 |
| 7 | BA1-Pro10 |
| 8 | BA2-WT |
| 9 | BA2-GP |
| 10 | BA2-G |
| 11 | BA2-P |
| 12 | BA2-AP10 |
| 13 | BA2-EAAAK4 |
| 14 | BA2-Pro10 |
| 15 | BA3-WT |
| 16 | BA3-GP |
| 17 | BA3-G |
| 18 | BA3-P |
| 19 | BA3-AP10 |
| 20 | BA3-EAAAK4 |
| 21 | BA3-Pro10 |
| 22 | BA1-WT |
| 23 | BA1-GP2 |
| 24 | BA1-GP4 |
| 25 | BA2-WT |
| 26 | BA2-GP2 |
| 27 | BA2-GP4 |
| 28 | BA1-GP8 |
| 29 | BA2-GP8 |
| 30 | BA3-WT |
| 31 | BA3-GP2 |
| 32 | BA3-GP4 |
| 33 | BA1- GP4 (+gly) |
| 34 | BA1-GP4 (+gly and –gly) |
| 35 | BA1-GP4 (-gly) |
| 36 | BA1-GP2 (+gly) |
| 37 | BA1-GP2 (-gly) |
| 38 | BA2-GP2 (+gly) |
| 39 | BA2-GP2 (-gly) |
| 40 | BA1-GP4 (+gly) |
| 41 | BA1-GP4 (-gly) |

**Supplementary Figure 5.** SDS-PAGE analysis of BpAbs with a peptide linker into one of heavy chains (**a**), with multiple GP linkers into both of heavy chains (**b**), before (+gly) and after (-gly) deglycosylation (**c**).

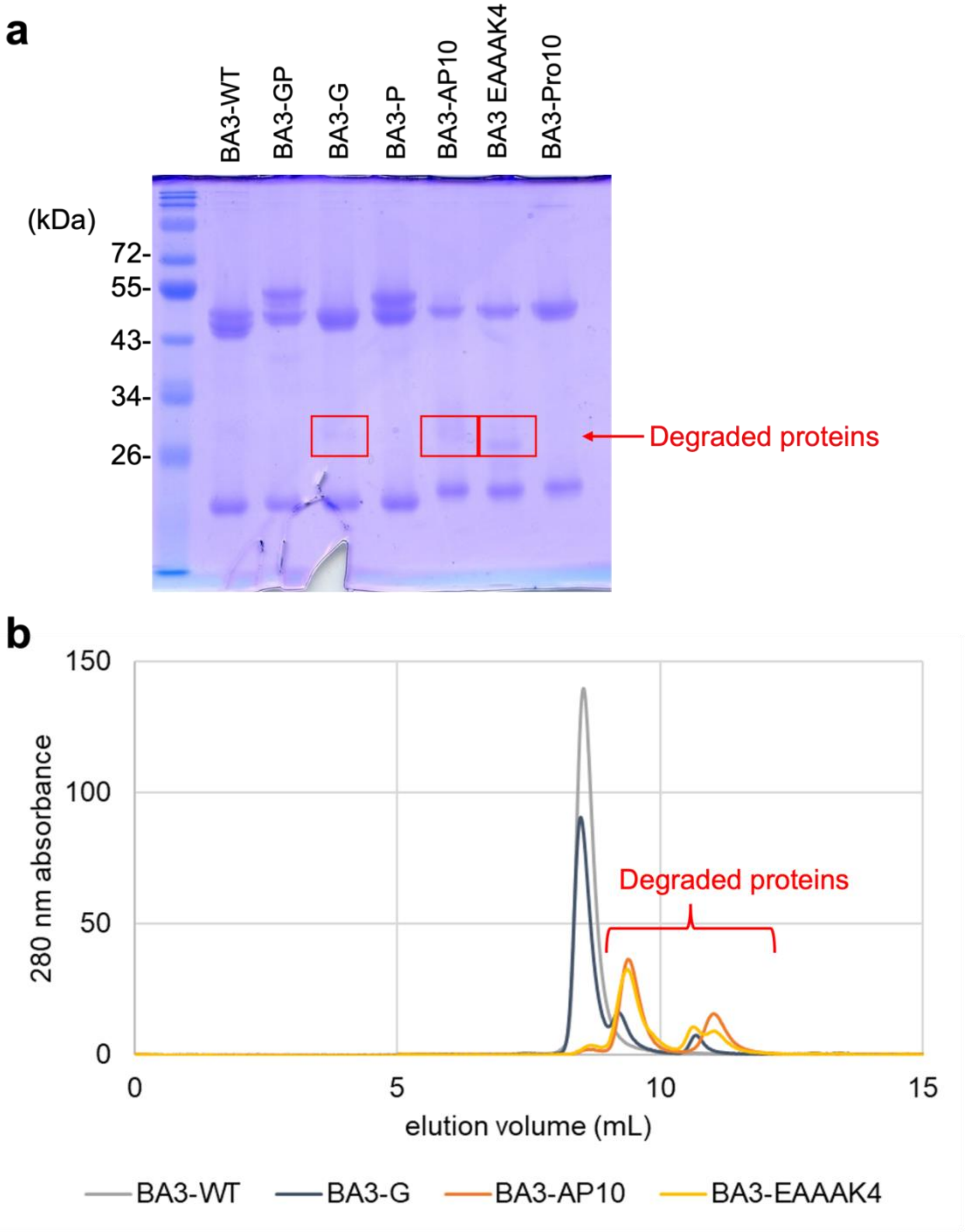


**Supplementary Figure 6.** Degradation of biparatopic antibodies stored at 4 °C, one month after purification. Analysis with SDS-PAGE (**a**) and with TSKgel G3000SWXL size-exclusion column (**b**).

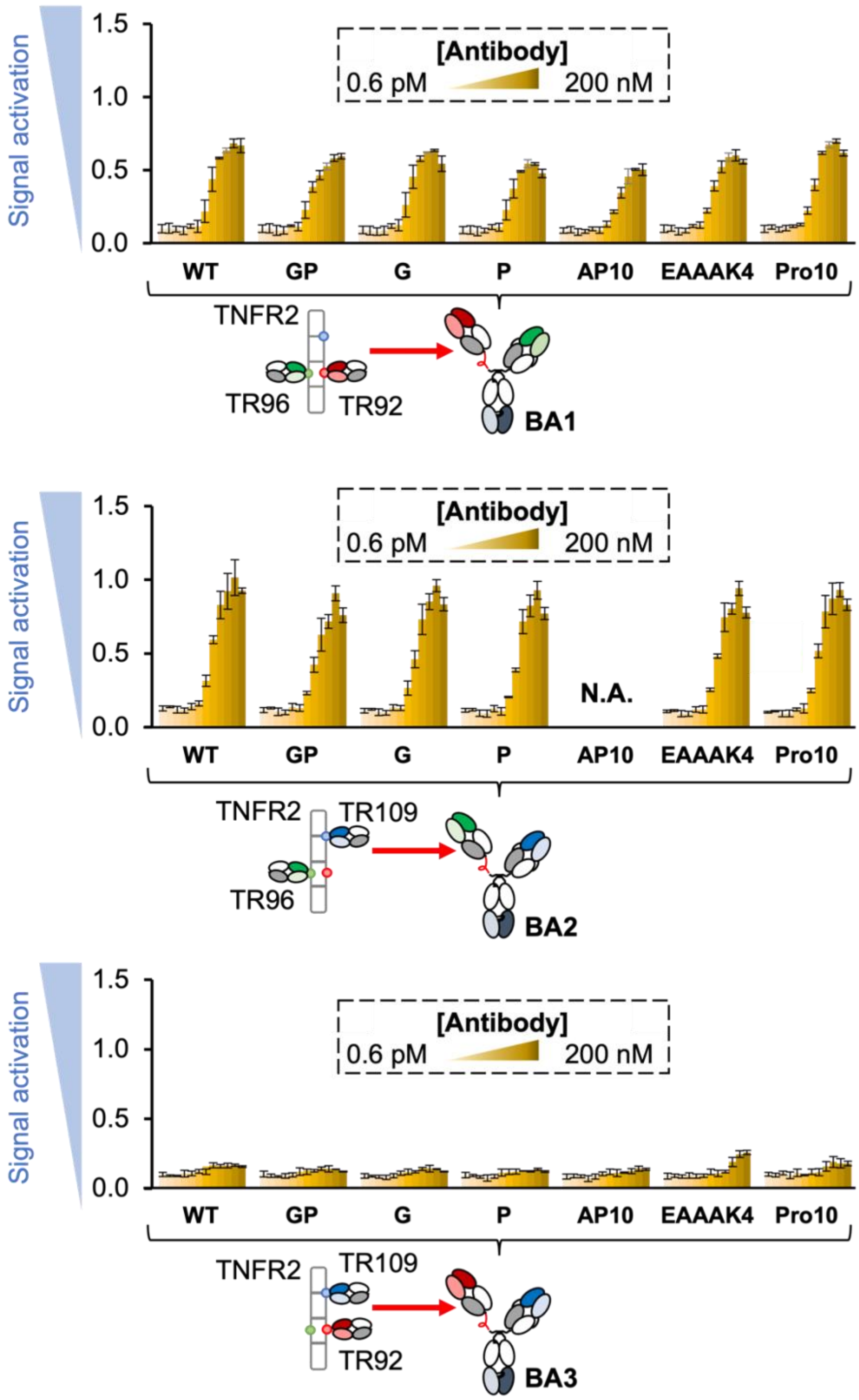


**Supplementary Figure 7.** Control of biological activities through linker insertion into the hinge region of one of heavy chains. Agonistic activities of BpAbs in a reporter gene assay are shown. BA2-AP10 was not analyzed due to degradation after purification. Values are shown with the standard deviation of three independent experiments.

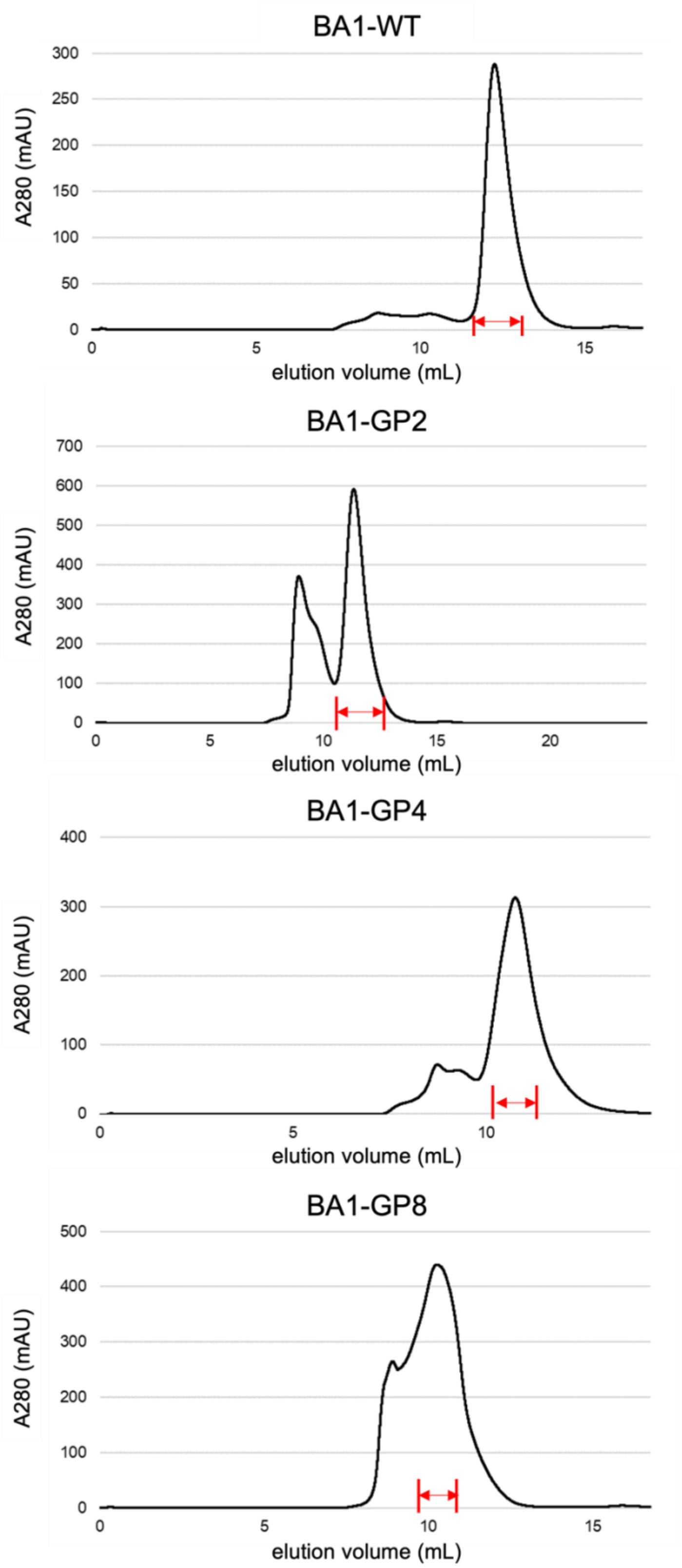


**Supplementary Figure 8.** Preparative size-exclusion chromatograms of BA1-WT, BA1-GP2, BA1-GP4 and BA1-GP8. The fractions with red arrows were collected.

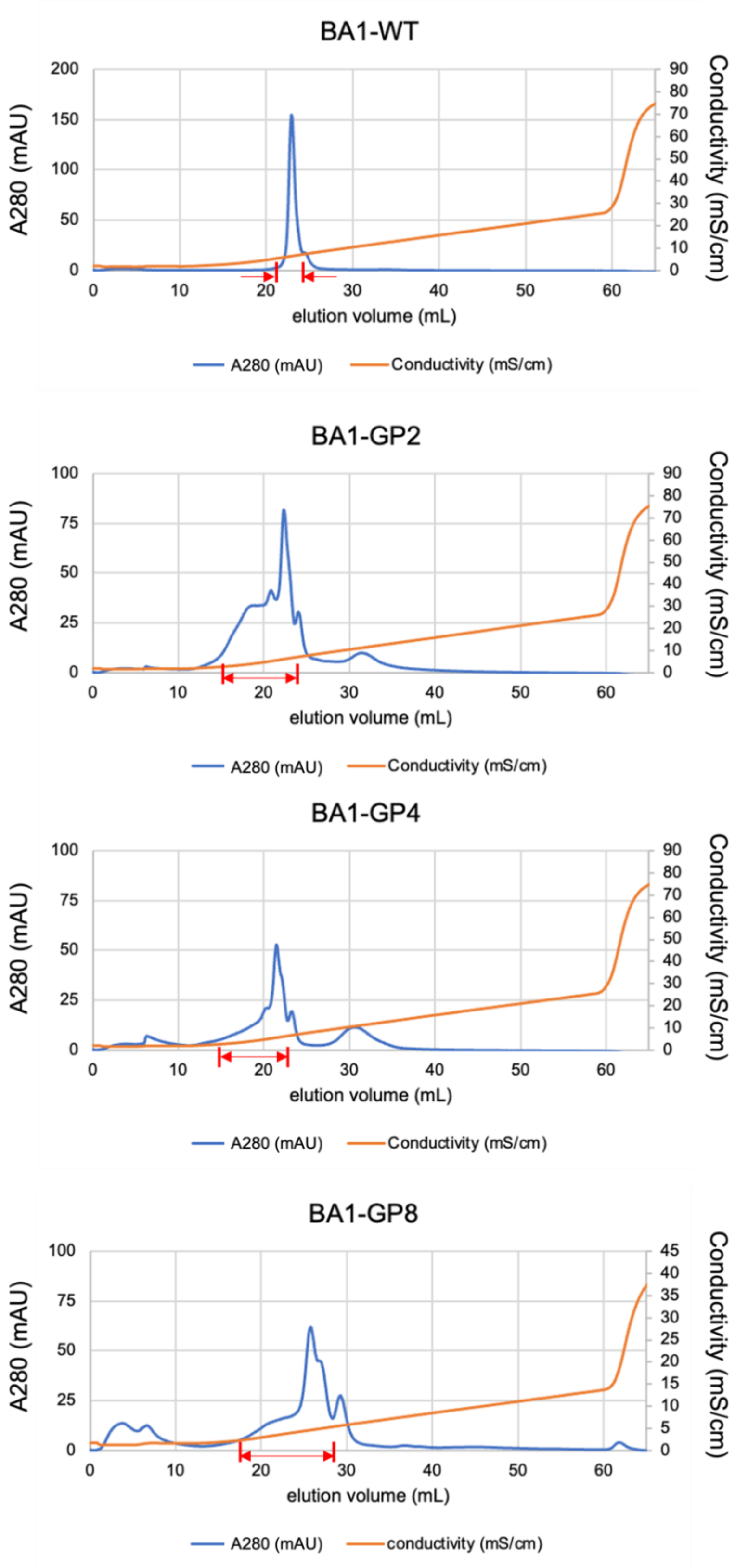


**Supplementary Figure 9.** Preparative cation-exchange chromatograms of BA1-WT, BA1-GP2, BA1-GP4 and BA1-GP8. The fractions with red arrows were collected.

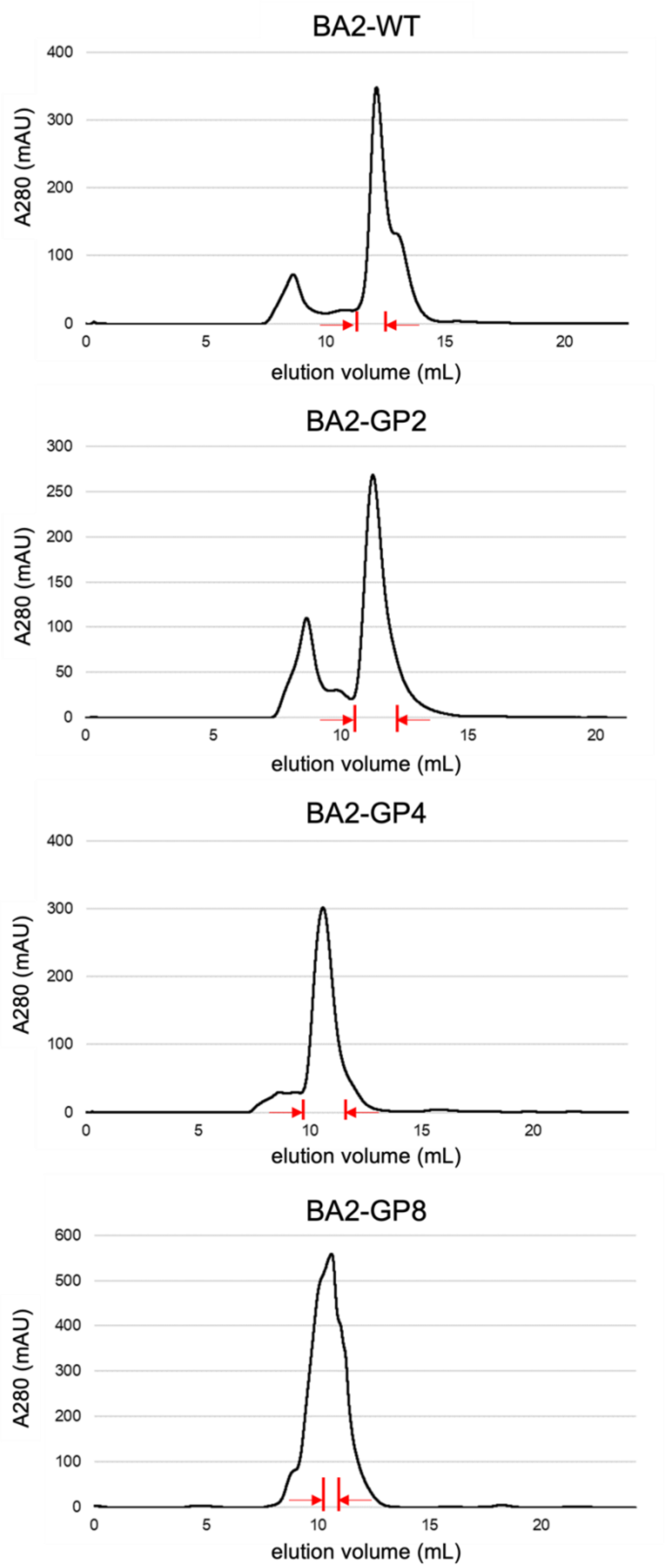


**Supplementary Figure 10.** Preparative size-exclusion chromatograms of BA2-WT, BA2-GP2, BA2-GP4 and BA2-GP8. The fractions with red arrows were collected.

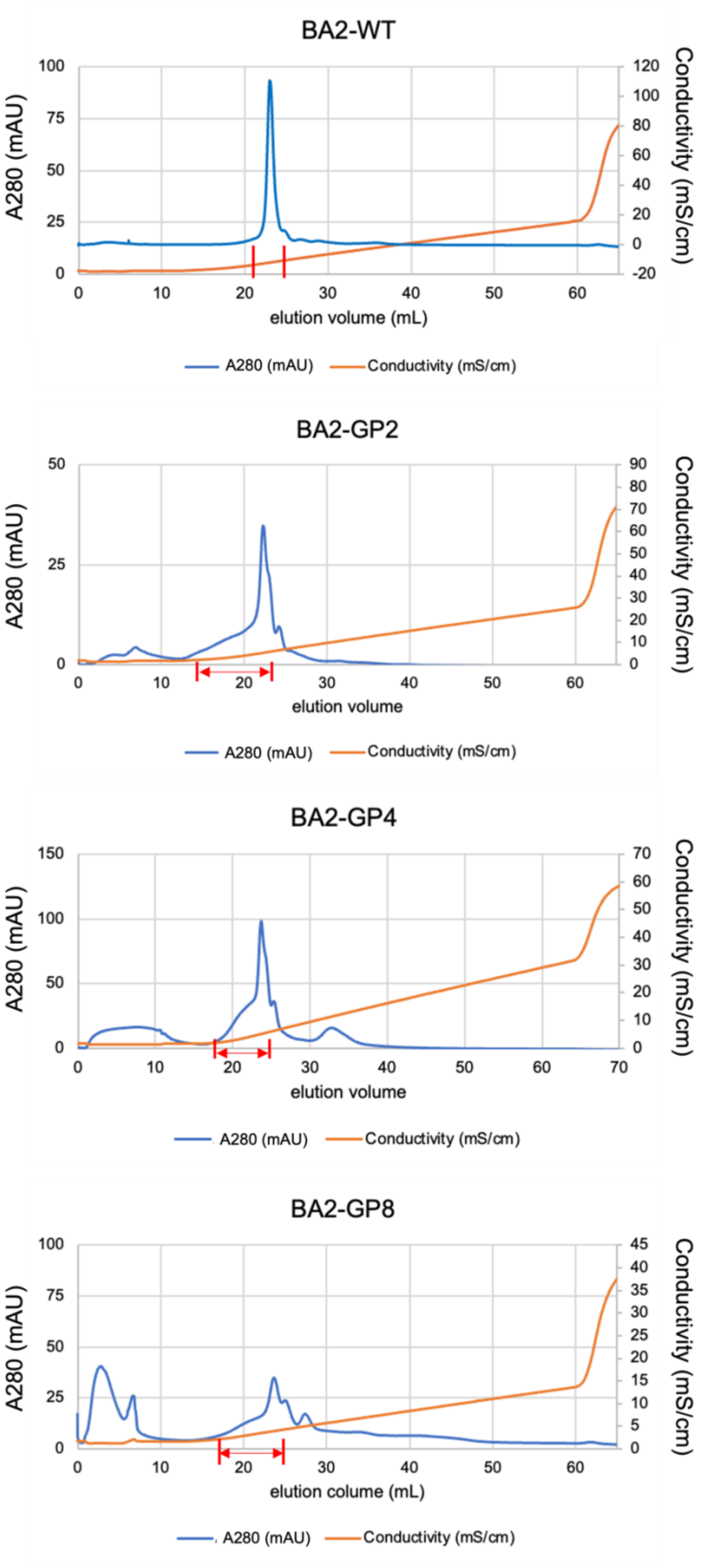


**Supplementary Figure 11.** Preparative cation-exchange chromatograms of BA2-WT, BA2-GP2, BA2-GP4 and BA2-GP8. The fractions with red arrows were collected.

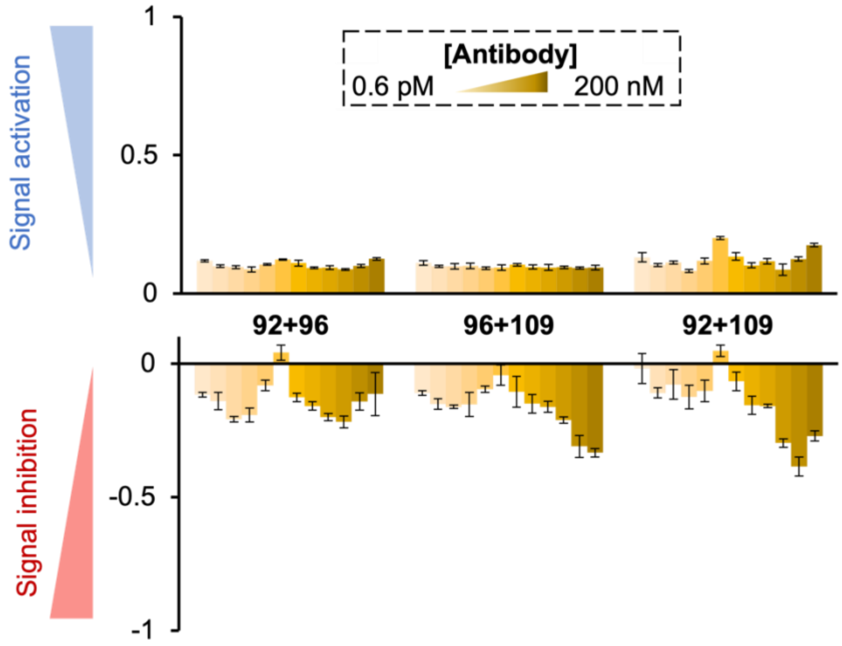


**Supplementary Figure 12.** Agonistic and antagonistic activities of the mixtures of two Fab proteins in a reporter gene assay. Values are shown with the standard deviation of three independent experiments.

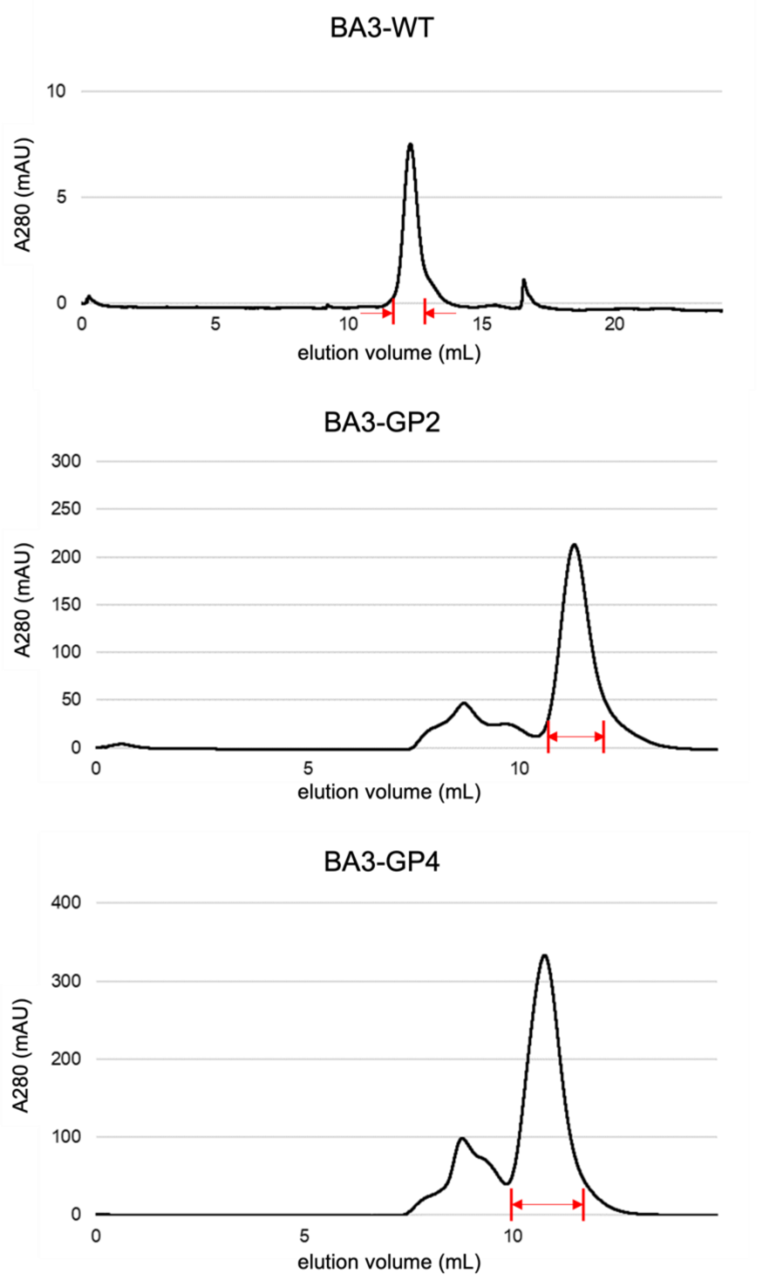


**Supplementary Figure 13.** Preparative size-exclusion chromatograms of BA3-WT, BA3-GP2 and BA3-GP4. The fractions with red arrows were collected.

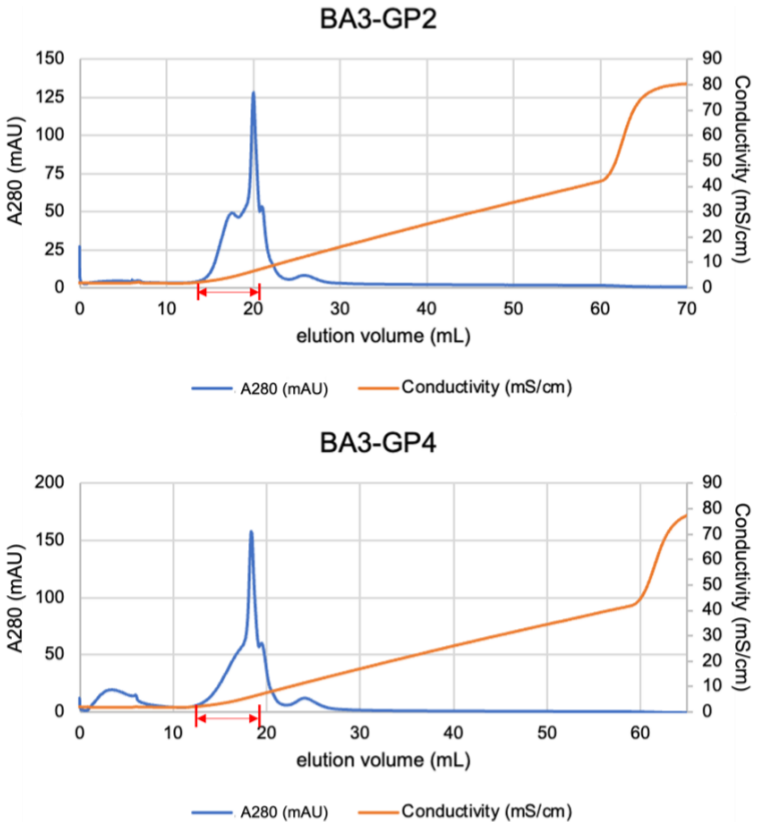


**Supplementary Figure 14.** Preparative cation-exchange chromatograms of BA3-GP2 and BA3-GP4. The fractions with red arrows were collected.

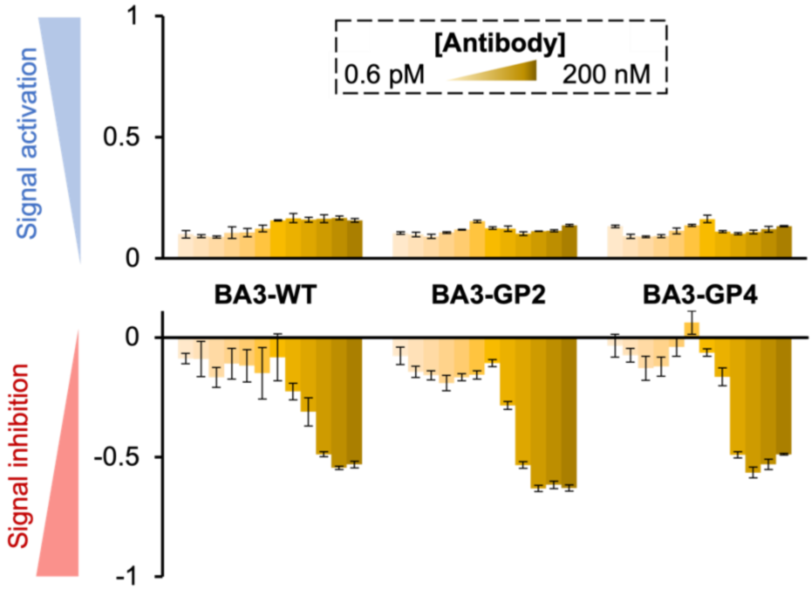


**Supplementary Figure 15.** Agonistic (upper) and antagonistic (lower) activities of BA3-WT, BA3 GP2 and BA3-GP4 in a reporter gene assay. Values are shown with the standard deviation of three independent experiments.

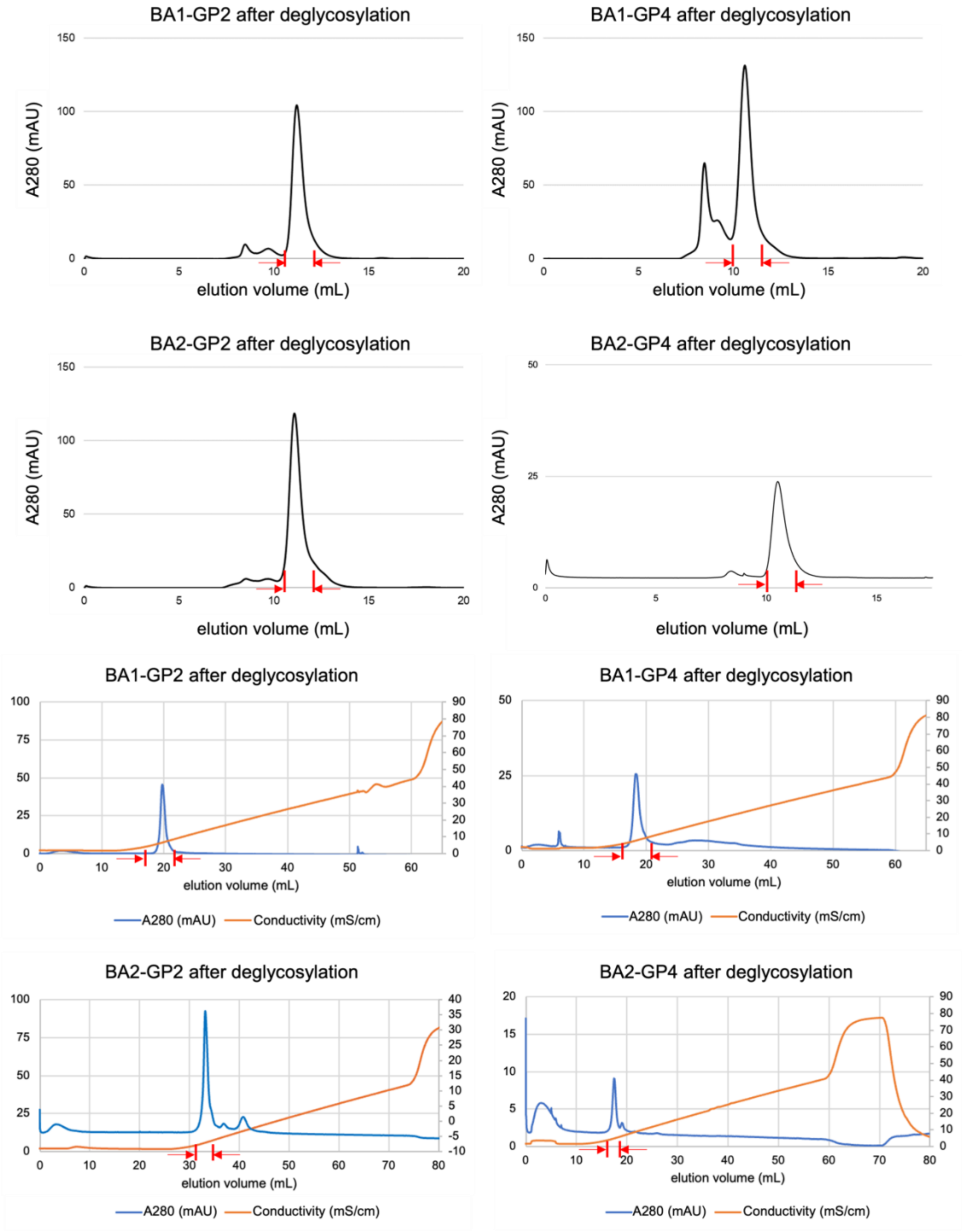


**Supplementary Figure 16.** Preparative size-exclusion chromatograms (upper four charts) and cation-exchange chromatograms (lower four charts) of BA1-GP2, BA1-GP4, BA2-GP2 and BA2-GP4 after deglycosylation. The fractions with red arrows were collected.

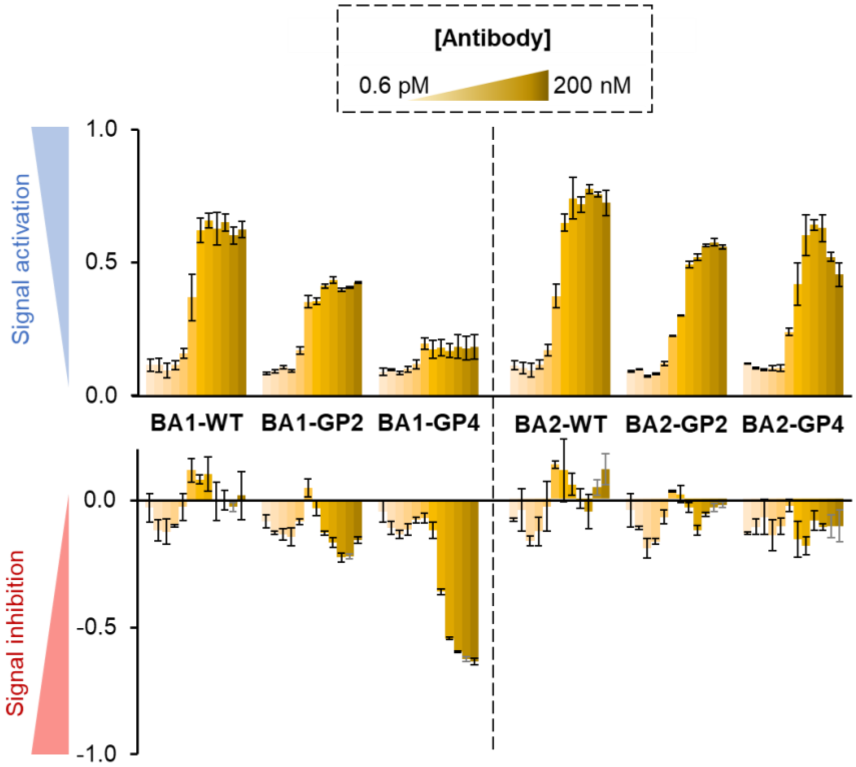


**Supplementary Figure 17.** Agonistic (upper) and antagonistic (lower) activities of BpAbs after deglycosylation in a reporter gene assay. Values are shown with the standard deviation of three independent experiments.

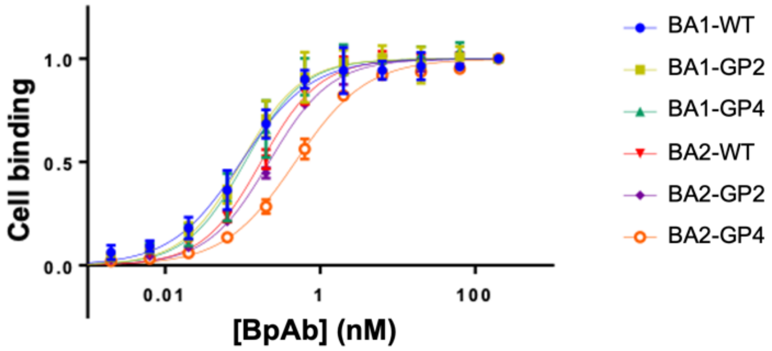


| | BA1-WT | BA1-GP2 | BA1-GP4 | BA2-WT | BA2-GP2 | BA2-GP4 |
|---|---|---|---|---|---|---|
| **log [EC50/nM]** | -1.021±0.041 | -1.015±0.035 | -0.971±0.038 | -0.763±0.026 | -0.659±0.017 | -0.321±0.018 |
| **EC50 (nM)** | 0.095 | 0.097 | 0.107 | 0.173 | 0.219 | 0.478 |

**Supplementary Figure 18.** Apparent binding affinity of BpAbs to TNFR2-expressing Ramos-Blue cells analyzed by using flow cytometry.

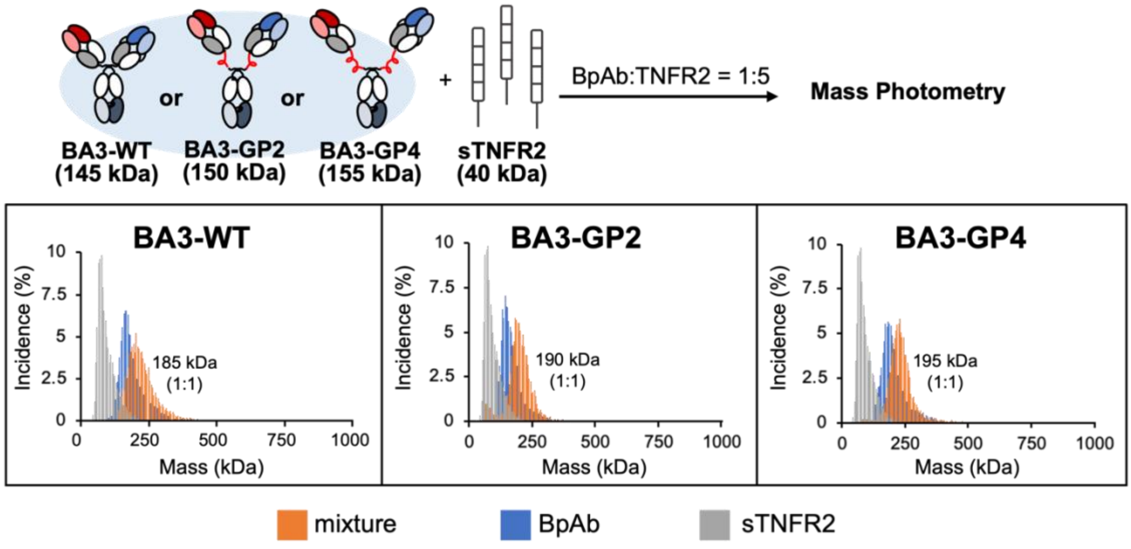


**Supplementary Figure 19.** Immunocomplex formed between BA3 and TNFR2. BpAbs were mixed with 5 eq. of sTNFR2.

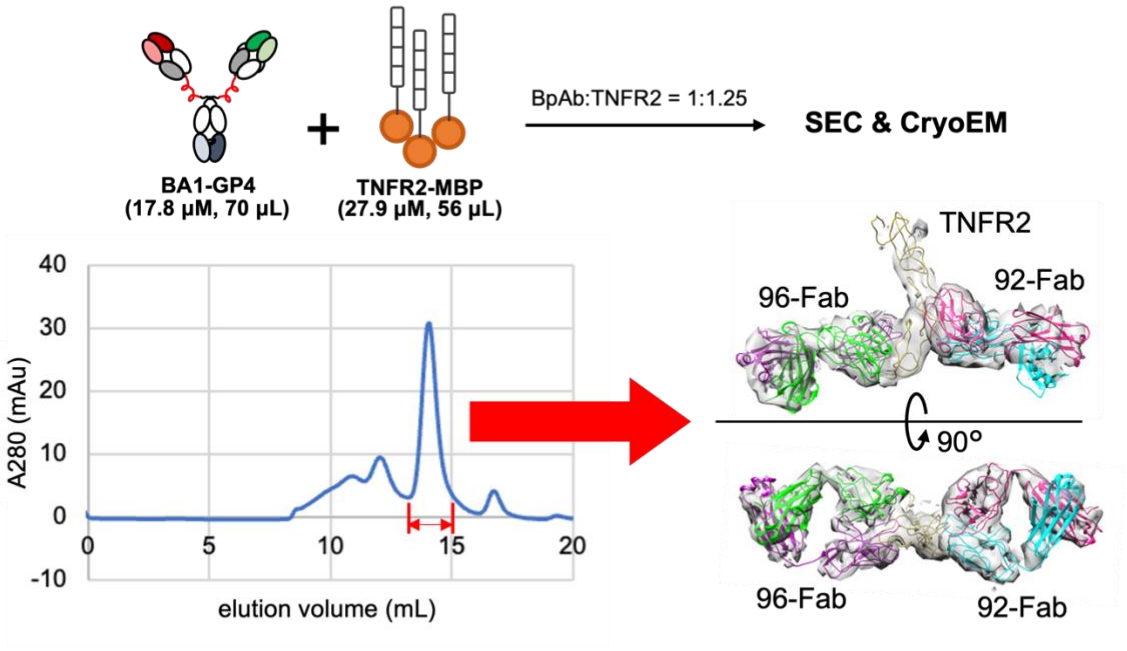


**Supplementary Figure 20.** Sample preparation for cryo-electron microscopy. Immunocomplex was collected by size-exclusion chromatography for cryo-electron microscopy.

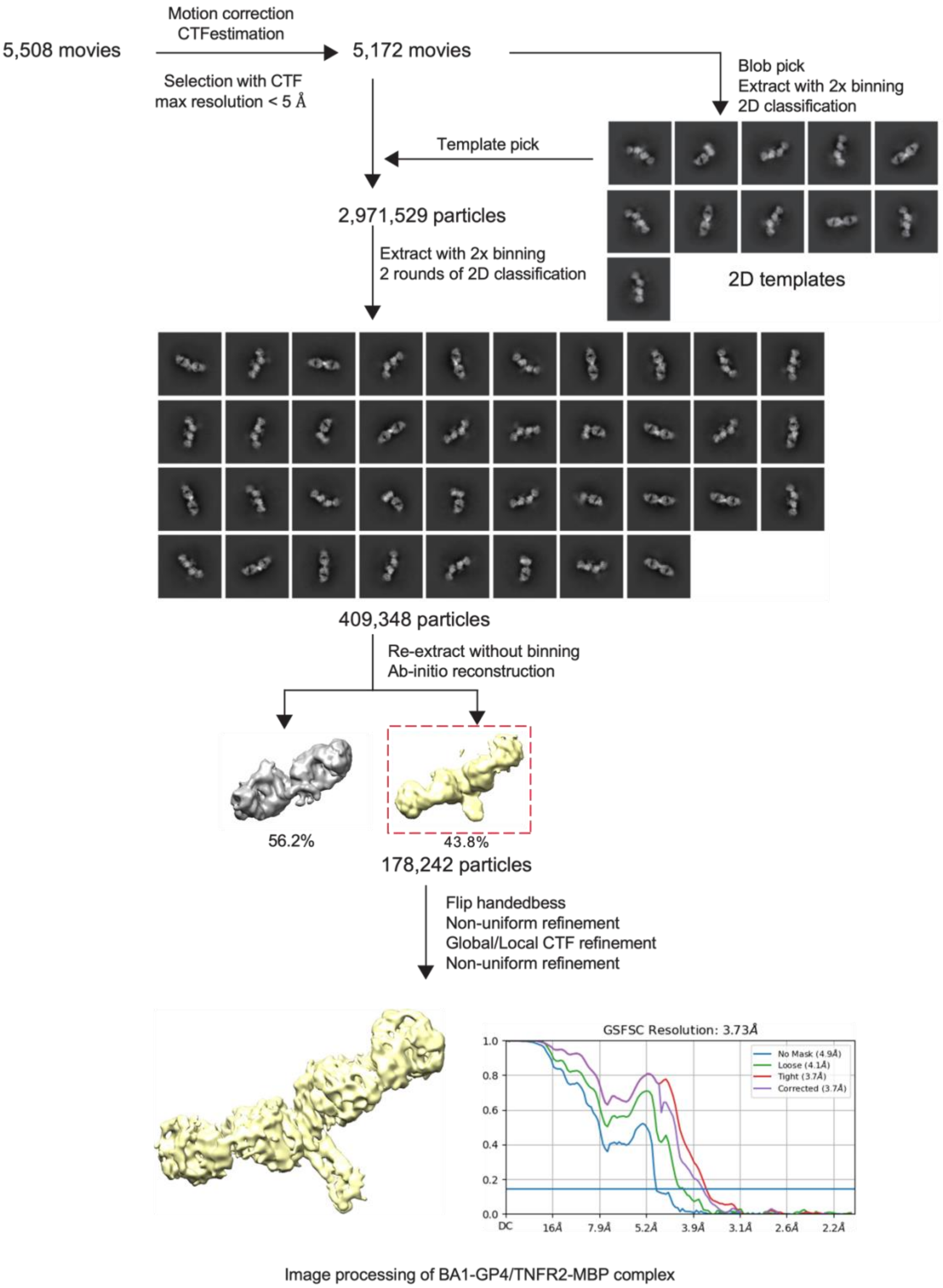


**Supplementary Figure 21.** Image processing of BA1-GP4/TNFR2-MBP complex.

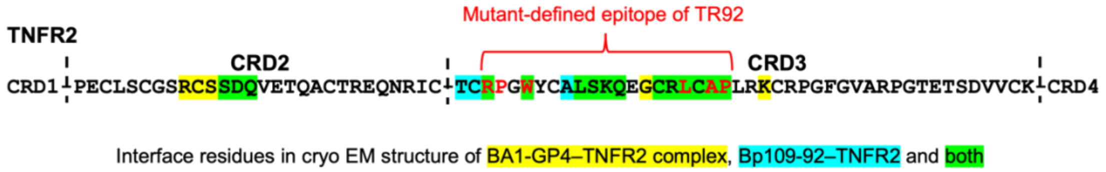


**Supplementary Figure 22.** Amino-acid residues of TNFR2 in the interface with 92-Fab from two different BpAbs. Interface residues are defined as Δ(buried surface area) > 0 calculated in PISA server. Interface residues that overlap in BA1-GP4–TNFR2 and Bp109-92–TNFR2 complexes are highlighted in green. Interface residues found only for BA1-GP4–TNFR2 or Bp109-92–TNFR2 are highlighted in yellow or cyan, respectively. Red texts indicate the core epitope residues defined by mutagenesis in the previous report (Akiba, H. *et al. Commun. Biol.* **6**, 987 (2023)).

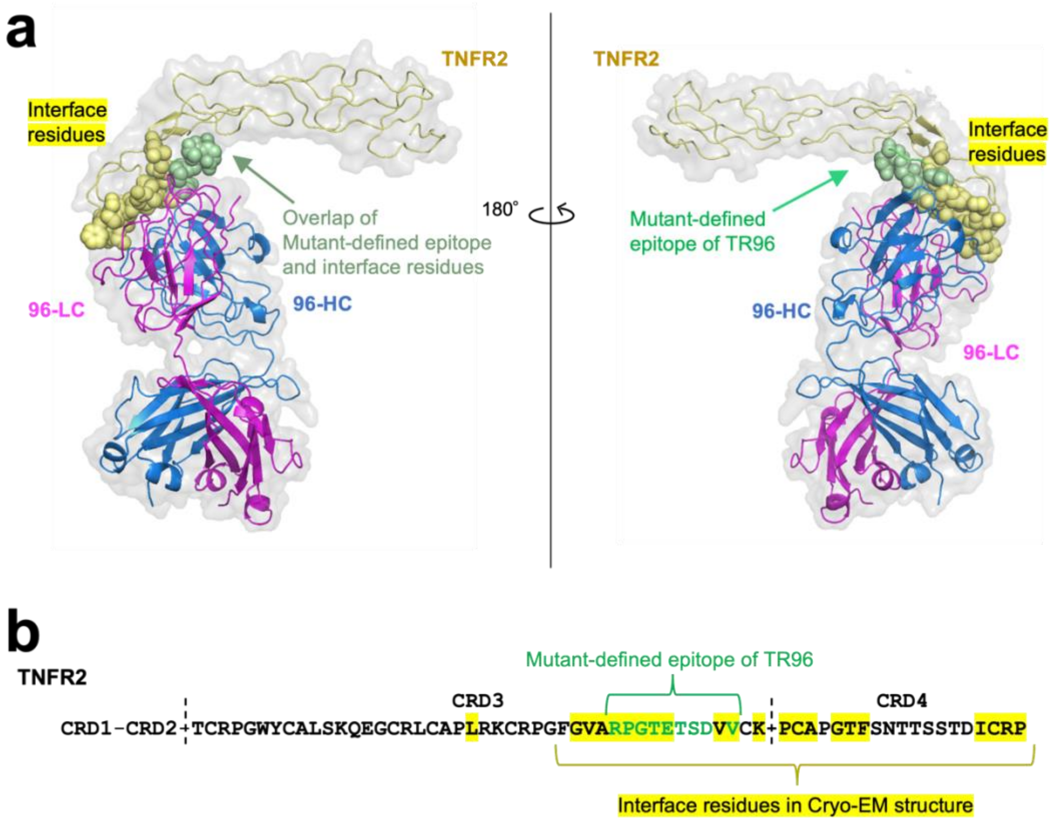


**Supplementary Figure 23.** The epitope of 96-Fab binding to TNFR2. (**a**) The epitope mapped on the complex structure of 96-Fab and TNFR2 from PDB ID 9LFL. Amino-acid residues of TNFR2 in the interface with 96-Fab are shown with spheres. Interface residues are defined as Δ(buried surface area) > 0 calculated in PISA server. TNFR2, 96-HC and 96-LC are respectively shown in yellow, blue and magenta. Epitope region found by mutagenesis in a previous study (Akiba, H. *et al. Commun. Biol.* **6**, 987 (2023)) are colored green. Because the mutagenesis was conducted by loop-level substitution, amino acid residues without direct contact are also included in this region. (**b**) Highlighted amino acid residues in (**a**).

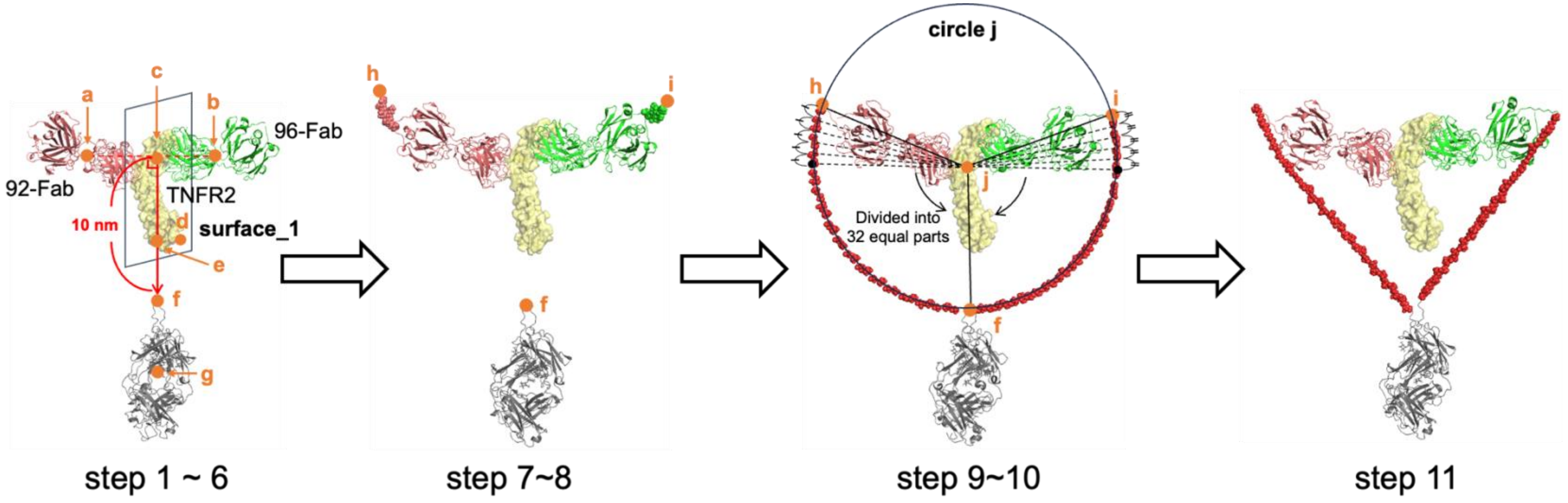


1. Centroids of all atoms in 92-Fab and 96-Fab are defined as **a** and **b**, respectively
2. **Surface_1** was defined as a plane that passes through the midpoint (**c**) of points **a** and **b**, with the line **ab** as the normal
3. Point **e** on the surface_1 was defined as the closest point to the N-terminal amino acid residue of TNFR2 (**d**)
4. Point **f** was defined on the line **ce**, located 10 nm away from **c**
5. N-terminal disulfide bond of Fc was positioned on **f**
6. Centroid of all amino acids in the Fc was defined as point **g** and positioned on the line **ce**
7. Missing C-terminal amino acid residues of two Fabs were inserted
8. C-termini of the heavy chains of 92-Fab and 96-Fab were defined as points **h** and **i**, respectively
9. A circle that passes through points **f**, **h**, and **i** was defined as circle **j**
10. Amino acid residues of the linkers were positioned at an equal interval along the circle **j**
11. Energy minimization was performed in MOE

**Supplementary Figure 24.** Strategy for building the initial structure of the complexes of BA1-GP2 and TNFR2.

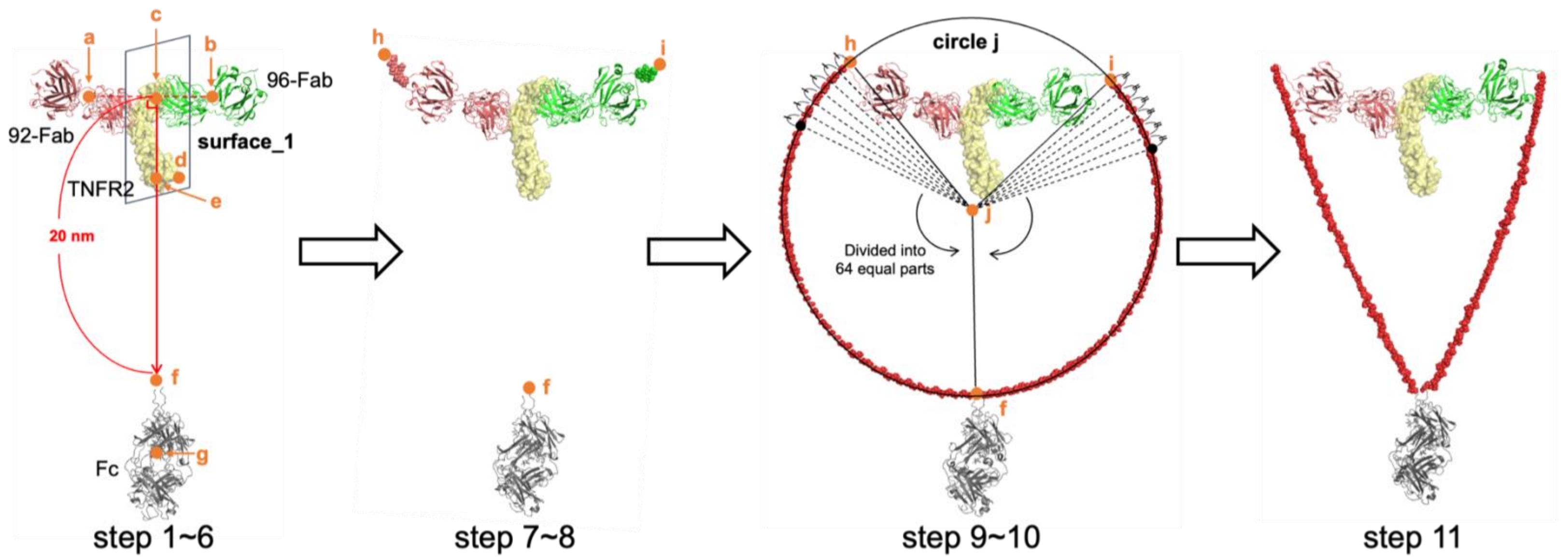


1. Centroids of all atoms in 92-Fab and 96-Fab are defined as **a** and **b**, respectively
2. **Surface_1** was defined as a plane that passes through the midpoint (**c**) of points **a** and **b**, with the line **ab** as the normal
3. Point **e** on the surface_1 was defined as the closest point to the N-terminal amino acid residue of TNFR2 (**d**)
4. Point **f** was defined on the line **ce**, located 20 nm away from **c**
5. N-terminal disulfide bond of Fc was positioned on **f**
6. Centroid of all amino acids in the Fc was defined as point **g** and positioned on the line **ce**
7. Missing C-terminal amino acid residues of two Fabs were inserted
8. C-termini of the heavy chains of 92-Fab and 96-Fab were defined as points **h** and **i**, respectively
9. A circle that passes through points **f**, **h**, and **i** was defined as circle **j**
10. Amino acid residues of the linkers were positioned at an equal interval along the circle **j**
11. Energy minimization was performed in MOE

**Supplementary Figure 25.** Strategy for building the initial structure of the complexes of BA1-GP4 and TNFR2.

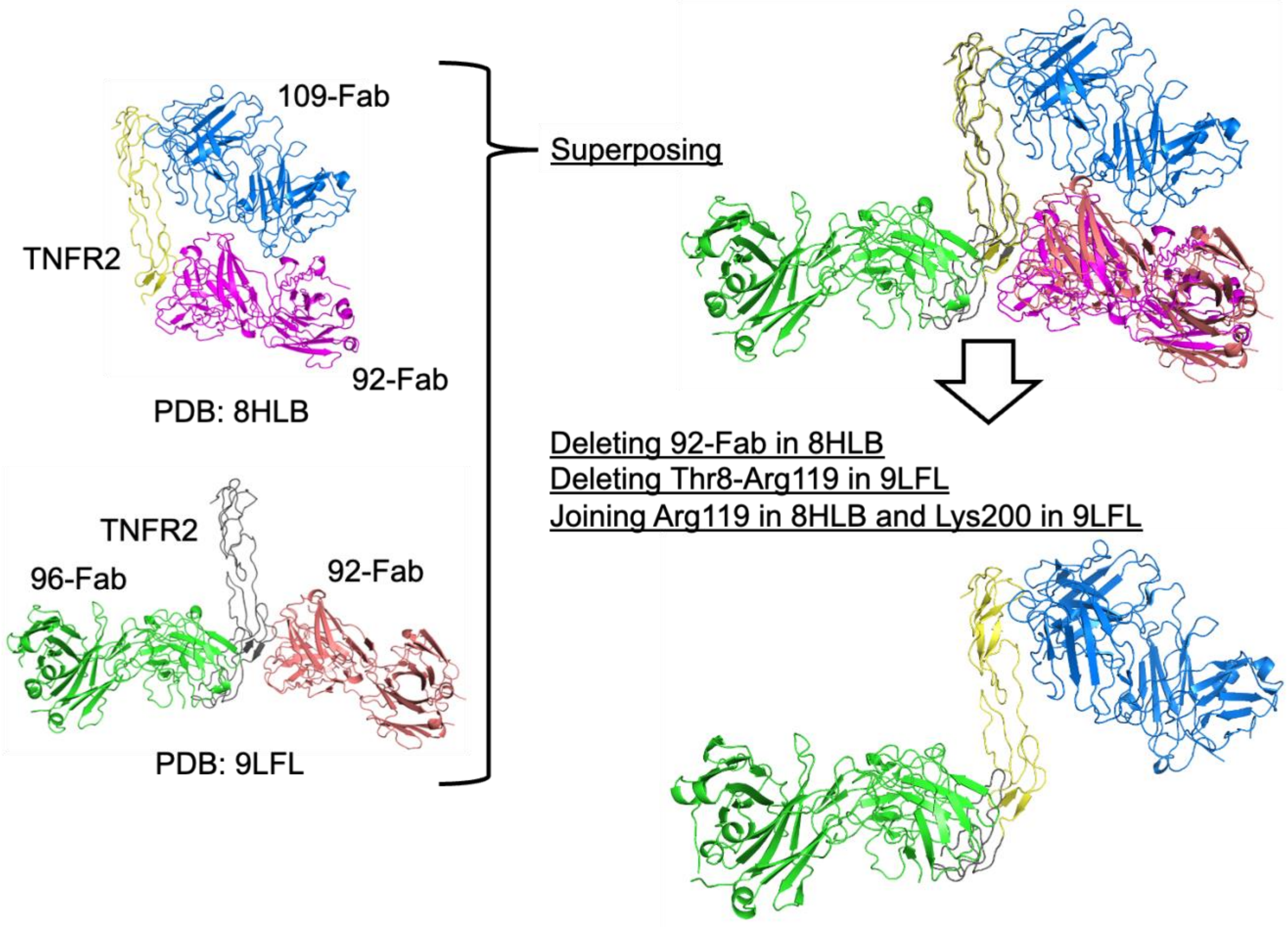


**Supplementary Figure 26.** Strategy for building ternary complex of 109-Fab, 96-Fab and TNFR2.

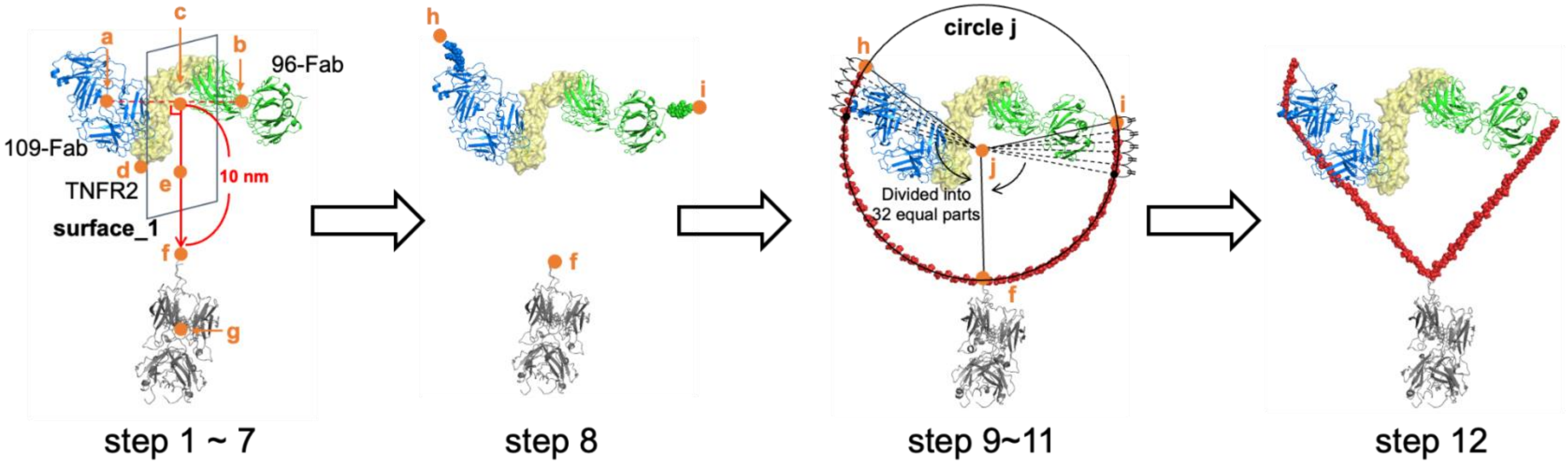


1. Centroids of all atoms in 109-Fab and 96-Fab are defined as **a** and **b**, respectively
2. **Surface_1** was defined as a plane that passes through the midpoint (**c**) of points **a** and **b**, with the line **ab** as the normal
3. Point **e** on the surface_1 was defined as the closest point to the N-terminal amino acid residue of TNFR2 (**d**)
4. Point **f** was defined on the line **ce**, located 10 nm away from **c**
5. N-terminal disulfide bond of Fc was positioned on **f**
6. Centroid of all amino acids in the Fc was defined as point **g** and positioned on the line **ce**
7. Missing C-terminal amino acid residues of two Fabs were inserted
8. C-termini of the heavy chains of 109-Fab and 96-Fab were defined as points **h** and **i**, respectively
9. A circle that passes through points **f**, **h**, and **i** was defined as circle **j**
10. Amino acid residues of the linkers were positioned at an equal interval along the circle **j**
11. Energy minimization was performed in MOE

**Supplementary Figure 27.** Strategy for building the initial structure of the complexes of BA2-GP2 and TNFR2.

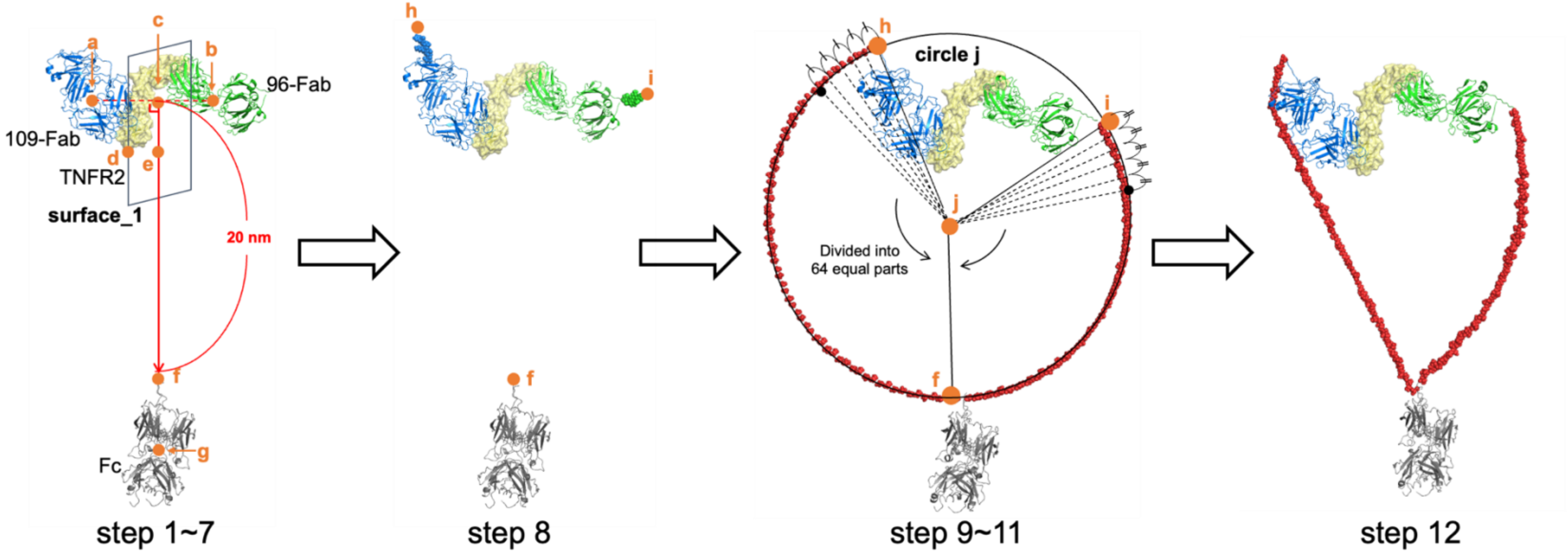


1. Centroids of all atoms in 109-Fab and 96-Fab are defined as **a** and **b**, respectively
2. **Surface_1** was defined as a plane that passes through the midpoint (**c**) of points **a** and **b**, with the line **ab** as the normal
3. Point **e** on the surface_1 was defined as the closest point to the N-terminal amino acid residue of TNFR2 (**d**)
4. Point **f** was defined on the line **ce**, located 20 nm away from **c**
5. N-terminal disulfide bond of Fc was positioned on **f**
6. Centroid of all amino acids in the Fc was defined as point **g** and positioned on the line **ce**
7. Missing C-terminal amino acid residues of two Fabs were inserted
8. C-termini of the heavy chains of 109-Fab and 96-Fab were defined as points **h** and **i**, respectively
9. A circle that passes through points **f**, **h**, and **i** was defined as circle **j**
10. Amino acid residues of the linkers were positioned at an equal interval along the circle **j**
11. Energy minimization was performed in MOE

**Supplementary Figure 28.** Strategy for building the initial structure of the complexes of BA2-GP4 and TNFR2.
